# Genome-Wide Association Study Reveals Influence of Cell-specific Gene Networks on Soybean Root System Architecture

**DOI:** 10.1101/2024.02.27.581071

**Authors:** Ying Sun, Charlotte Miller, Ashish B. Rajurkar, Ryan C. Lynch, Anthony Alyward, Ling Zhang, Marieken Shaner, Charles D. Copeland, Heng Ye, Henry T. Nguyen, Wolfgang Busch, Todd P. Michael

**Affiliations:** The Plant Molecular and Cellular Biology Laboratory, The Salk Institute for Biological Studies, La Jolla, CA 92037, USA; Division of Plant Sciences and National Center for Soybean Biotechnology, University of Missouri-Columbia, Columbia, MO, 65211, USA; Department of Biology, San Diego State University, San Diego, CA 92182; School of Global Policy and Strategy, University of California, La Jolla, CA 92093

**Keywords:** genome-wide association study (GWAS), Soybean, Root system architecture (RSA), metaphloem, endodermis, lateral root length

## Abstract

Root system architecture (RSA) describes the shape and arrangement of a plant’s roots in the soil including the angle, rate of growth, and type of individual roots, which facilitates the uptake of nutrients and water. In crop improvement efforts, RSA has been less well studied due to the technical challenges associated with phenotyping roots as well as a focus on above-ground traits such as yield. We developed a gel-based root phenotyping system called *RADICYL* (Root Architecture 3D Cylinder), which is a non-invasive, high-throughput approach that enabled us to measure 15 RSA traits. We leveraged *RADICYL* to perform a comprehensive genome-wide association study (GWAS) with a panel of 371 diverse soybean elite lines, cultivars, landraces, and closely related species to identify gene networks underlying RSA. We identified 54 significant single nucleotide polymorphisms (SNPs) in our GWAS, some of which were shared across multiple RSA traits while others were specific to a given trait. We generated a single cell atlas of the soybean root using single nuclei RNA sequencing (snRNAseq) to explore the associated genes in the context of root tissues. Using gene co-expression network (GCN) analyses applied to RNA-seq of soybean root tissues, we identified network-level associations of genes predominantly expressed in endodermis with root width, and of those expressed in metaphloem with lateral root length. Our results suggest that pathways active in the endodermis and metaphloem cell-types influence soybean root system architecture.

## Introduction

The spatial distribution of roots, Root System Architecture (RSA), is a key determinant for the ability of roots to capture nutrients and water from the soil environment, which strongly influences plant fitness and yield. RSA arises through root growth, root growth direction, and root branching (Slovak et al. 2016), which are influenced by both genetic and environmental factors (Lynch 2022). While the root system in dicots consists of a single primary root that can develop several orders of lateral roots, the monocot root system contains primary and seminal roots as well as shoot-borne roots and lateral roots. In both cases, the lateral branching is an important determinant of RSA and is based on the post-embryonic development of lateral roots from the pericycle (Parizot et al. 2008; Slovak et al. 2016), a tissue layer between the central vascular cylinder and endodermis. Lateral root initiation involves a population of founder cells in the pericycle layer that are specified through an auxin-dependent process (De Smet et al. 2007) and once activated, start to divide and form a lateral root primordium that can develop into a lateral root. The location of lateral root initiation sites is highly regulated and has been shown in the model species *Arabidopsis thaliana* to depend on an oscillatory clock-like process (Wachsman et al. 2020). Additionally, many other signaling mechanisms involving receptor-like kinases (RLKs) and various factors contribute to lateral root development (Rodriguez-Villalon et al. 2015; Jourquin, Fukaki, and Beeckman 2020; Ou, Kui, and Li 2021).

While genes and molecular processes involved in RSA have been extensively studied in *Arabidopsis*, they are understudied in crop species. One of the most important crop species is soybean (*Glycine max*), which ranks as the fourth largest crop globally. Soybean seeds contain high protein and edible oil levels and are used for human consumption, animal feed, and oil production (Guo et al. 2022; Zhao et al. 2017). Several genome-wide association studies (GWAS) have been conducted to study RSA in soybean and many of these studies focused on RSA data that was obtained by growing roots in environments restricting their growth to two-dimensions (2D). Two of the studies utilized pouch and wick systems based on growing them on moistened blue paper (Falk et al. 2020, Chandnani et al. 2023). While RSA could be accurately quantified in these studies, the 2D root growth is very far removed from environments found in the field. Other studies used restricted soil-based root systems, soil grown roots either in seedling cone systems, rhizoboxes or PVC pipes (8 cm diameter and 35 cm height). Images of these space restricted root systems were obtained in a 2D way either by a flatbed scanner or by a camera taking an image of the flat surface of the rhizobox (Prince et al. 2019, Seck et al. 2020, Mandozai et al. 2020). Finally, a GWAS was conducted on crown roots of field grown soybeans, providing trait associations of the uppermost root system parts at the end of a field season (Dhanpal et al. 2020). Overall, these studies did not address soybean root systems that grow unconstrained in a three-dimensional (3D) environment or quantify developmental traits such as growth rate or the angles of the developing taproot vs. lateral roots. Moreover, despite previous research, there is still a lack of mechanistic insights into the formation of soybean RSA, especially in the roles of various genes that collectively contribute to significant effects.

One promising avenue for prioritizing candidate genes found in GWAS is to employ gene regulatory networks (GRN). GRN have contributed to several phenotypic discoveries including developmental patterns in fruit flies and sea urchins, the circadian rhythm of plants, flowering time regulation, and plant responses to abiotic stress (Tarsis et al. 2022; Imaizumi 2010; Sun et al. 2022). The increasing volume of expression data has popularized gene co-expression network (GCN) approaches as a proxy for GRN. Several strategies have been developed to construct GCNs that depict mRNA as nodes, co-expression relationships as edges, and co-expressed modules as connected components (Tantardini et al. 2019; Huynh-Thu et al. 2010; Cliff et al. 2019; Moerman et al. 2019; R. Zheng et al. 2019; Sun and Dinneny 2018). Weighted gene co-expression network analysis (WGCNA) has been applied to predict tissue-specific networks, identify networks related to lateral root and nodule formation, and identify the regulatory components of flooding tolerance (Jhan et al. 2023; Smita et al. 2020; Juexin Wang et al. 2019). GCN approaches using RNA-seq have been successfully applied to various crops, including soybean (Azam et al. 2023; Yao et al. 2023; Gao et al. 2018). Identifying network components representing sub-networks connected by central connecting genes could offer valuable insights into their regulatory roles for both plant and other development-related traits (Ko and Brandizzi 2020; X. Zhu, Duren, and Wong 2021). To gain greater insight into physiological mechanisms, single-cell and single-nuclei RNA-seq (scRNA-seq/snRNA-seq) could be supplemented to unveil the function of network components with cell-level resolution (Jia et al. 2022; Jagadeesh et al. 2022). In plants, these technologies have already uncovered novel developmental phenotypes, defined spatial and temporal patterns during biotic stress, facilitated comparisons of common cell-types in *Arabidopsis* and crops, and identified specialized cell-types and specific gene expression patterns (Dorrity et al. 2021; Shahan et al. 2022; J. Zhu et al. 2023; Apelt et al. 2022; Yılmaz et al. 2023; Song et al. 2020; Guillotin et al. 2023; Shahan, Nolan, and Benfey 2021). In soybeans, scRNA-Seq has also played a crucial role in unraveling the rhizobium-legume symbiosis by classifying major cell-types in both the root and root nodules (Liu et al. 2023).

Here we integrate GWAS, GCN, and snRNA-seq analysis to uncover the genetic regulation that contributes to shaping RSA in soybeans. In our efforts to provide an unrestricted root growth environment, we have developed a high-throughput 3D imaging system called **R**oot **A**rchitecture 3**D I**maging **Cyl**inder (*RADICYL*), which is based on the concept of growing seedlings unimpededly in cylinders filled with transparent gel media (Clark et al. 2011; Iyer-Pascuzzi et al. 2010). *RADICYL* provided advancements over previous methods in several aspects including size, hardware, the imaging camera, and system throughput. We developed a deep learning image segmentation approach that is capable of accurately quantifying RSA traits from the resulting images in a non-supervised manner. Employing these tools, we successfully quantified root traits in 371 diverse soybean varieties. Subsequently, we used publicly available whole genome resequencing data (Valliyodan et al. 2021) to conduct GWAS using multi-locus models (Jiabo Wang and Zhang 2021). We generated a single cell expression atlas of the soybean root to construct a gene co-expression network (GCN) that identifies sub-networks containing GWAS candidate genes. Examining these gene sets, we identified a subset of genes that are (a) highly expressed in endodermis and (b) proximal to SNPs associated with root width. Likewise, another gene subset associated metaphloem with lateral root length. These findings suggest key biological processes in these cell-types that shape variation in soybean RSA.

## Results

### RADICYL: A high-throughput phenotyping platform for screening Root System Architecture (RSA) traits

Root system architecture (RSA) is a highly composited trait that is best quantified using an imaging platform capable of capturing the root system in its natural three-dimensional (3D) state. We developed a high-throughput phenotyping platform, ***R****oot **A**rchitecture 3**D I**maging **Cyl**inder* (*RADICYL*) to achieve the necessary throughput required for screening such traits in the context of natural variation studies. In developing *RADICYL* we used the principles laid out by a previously published imaging method that makes use of gel-filled cylinders within which rice roots could grow unimpededly for approximately two weeks (Clark et al. 2011; Iyer-Pascuzzi et al. 2010). We enhanced throughput by decreasing the quantity of the gel medium required and utilized smaller polystyrene containers, instead of expensive and heavy glass containers (Clark et al. 2011; Iyer-Pascuzzi et al. 2010). Using *RADICYL,* a single person can screen 75 cylinders per hour. Within this time, the user was able to acquire a rotational image series consisting of 72 images at 5° intervals, giving a total of 5,400 captured images (Supplementary Fig. 1). We developed a deep-learning-based image processing pipeline to process the large number of images collected. In this process, the images were trimmed to a size of 990 x 860 pixels to center the plant within the cylinder and eliminate pixels outside of the cylinder area (Supplementary Fig. 2a). We then utilized a trained model obtained using the convolutional neural network (CNN)-based semantic segmentation architecture UNet++, to segment primary roots and lateral roots in each image. Following this, we skeletonized the roots in each segmented image and measured a total of 15 traits. The majority of the traits were directly computed based on the image (width, depth, convex hull area, biomass, total root length, primary root length, lateral root length, lateral root tip depth, primary root tip depth, all tip depth, vertical angles), while 3 were compound traits (SDx, SDy, and SDx/SDy) and 1 physical mass trait (dry root biomass) was measured after imaging on dried roots (Supplementary Fig. 2b, Table 1). We also implemented an efficient quality control (QC) step to discard samples with poor germination or other quality issues and to achieve higher fidelity. Overall, using the *RADICYL* imaging setup and the image processing workflow described, we were able to capture high-quality data elucidating a wide range of RSA traits in soybeans.

### Early RSA traits cylinder-grown soybean seedlings are heritable and display notable natural variation

We screened the 15 RSA parameters described above across a diverse set of 371 USDA soybean accessions using the *RADICYL* phenotyping platform. This collection of germplasm comprises a high level of genetic diversity with genotypes collected from more than 20 countries and across a range of maturity groups. Using the *RADICYL* system to evaluate the 15 traits across these genotypes uncovered a high level of phenotypic variation (Supplementary Fig. 3a, Table 2). We performed a correlation analysis to explore the relationship between the early RSA traits screened and identified traits with the highest positive correlation to be convex hull area, biomass area, lateral root length, and total root length, which showed an R^2^ of 0.9 (Supplementary Fig. 3b). The overall orientation of the root in the X direction (SDx) is strongly positive (R^2^ = 0.9) correlated with the width of the root system. Similarly, the orientation in the Y direction (SDy) is positively correlated (R^2^ = 0.7) with the depth of the root system. Early-stage root biomass was predominantly predicted by primary root length and lateral root length, and dry root biomass showed positive correlations (R^2^ = 0.4-0.5) with both of these root length traits.

Additionally, the dry root biomass displayed a strong positive correlation (R^2^ = 0.6) with biomass area, indicating that the root system captured in pixels is predictive of the actual physical mass of the early seedling stage root structures. The identification of relatively high positive correlations between traits suggests that these traits may measure similar aspects of the root and despite employing different measurement approaches, there might be a commonality in capturing comparable attributes.

We generated a Principal Component Analysis (PCA) plot based on the value of each trait across all accessions to visually represent the distribution and further explore the twelve directly computed traits (Figure 1a). The results demonstrated a clear separation of these traits along two distinct axes: PCA1 and PCA2, which collectively accounted for over 50.7% of the variance. The principal components separated the traits into mostly two clusters: the first cluster comprised traits including lateral root tip depth, primary root tip depth, all tip depth, primary root length, and depth, which we collectively referred to as “depth traits.” The second cluster contained traits such as width, lateral root length, biomass area, total root length, and convex hull, which we collectively referred to as “size and shape traits.” We next identified the accessions with the most significant phenotypic variability across the quantified traits by ranking the accessions in descending order of overall variation across all traits (Supplementary Fig. 4) and summarizing the top four accessions in a radial plot (Figure 1b). The accessions PI548540, PI548667, PI597464, and PI603458A underscored that the most significant differences in root architecture traits are associated with the number, length, and angle of the lateral roots (Figure 1c). Our data suggest that these accessions constitute the most variable subset within our dataset, highlighting the potential for identifying and selecting accessions using our methods.

**Figure 1.**
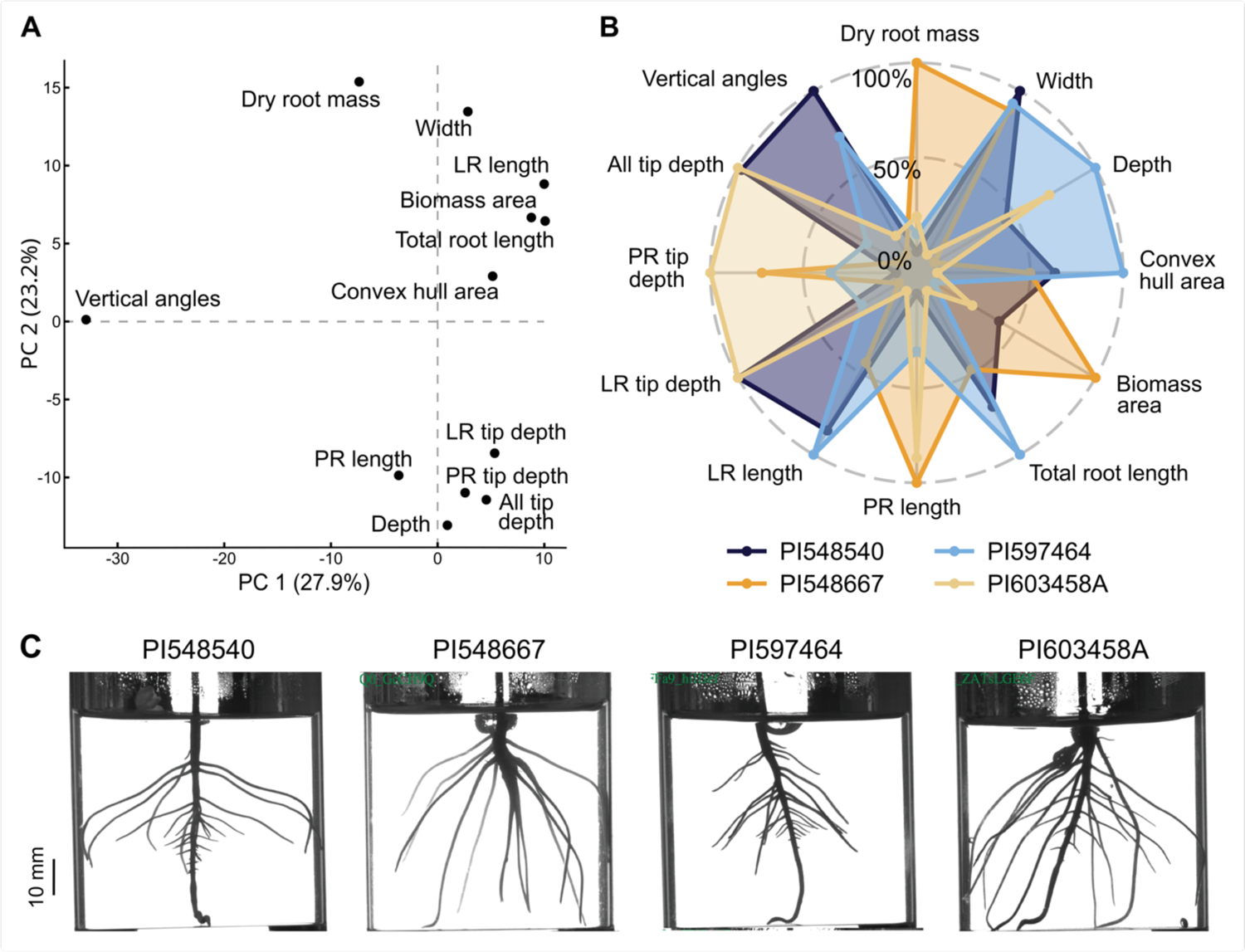
*RADICYL* captures a substantial range of variation in root system architecture (RSA) through the characterization of 12 directly computed trait measurements. A) Principal Component Analysis (PCA) plot of directly computed traits using phenotype data for all samples. B) A radial plot showcasing the top four accessions characterized by the highest cumulative variance across traits that exhibit the most significant deviations from the mean. C) Representative image of 6-day-old seedlings corresponding to the cylinder data presented in panel B, illustrating the phenotypic variation among accessions.

We calculated broad-sense heritability (BSH) from all the accessions to gain an understanding of how much of the phenotypic variation observed can be accounted for by genetic variation. BSH values ranged from 13% to 32%, with lateral root length showing the lowest BSH and dry biomass showing the highest. The identification of heritable variation for these traits is promising and emphasizes the inherent potential of the population to capture a substantial proportion of variability in RSA traits.

### Genome-Wide Association Studies (GWAS) pinpoint genetic associations and hotspots for soybean root architecture

We set out to use Genome Wide Association Studies (GWAS) to uncover the genetic regulation underlying the trait variation observed across the 371 soybean genotypes screened. The samples for our study encompassed diverse accessions, including cultivars and landraces from China (222), USA (52), Korea (36), Japan (23) and Russia (11) (Figure 2a). Genotypes across the 371 phenotyped accessions were called from publicly available whole genome Illumina sequencing data, yielding 4,815,704 high-quality single nucleotide polymorphisms (SNPs). The distribution of SNPs across the genome varied from 0 to 14,774 per megabase (Mb) (Supplementary Fig. 5). A PCA of accession genotypes found 34.83% of variance was explained by two PCs, which distinguished a cluster of cultivar type accessions from landrace type accessions (Figure 2b). The top two PCs did not separate accessions by maturity group, and the distribution of geographic origins was explained by the correlation of geography with accession type (most cultivars from the USA, most landraces from Asia) (Supplementary Fig. 6).

**Figure 2.**
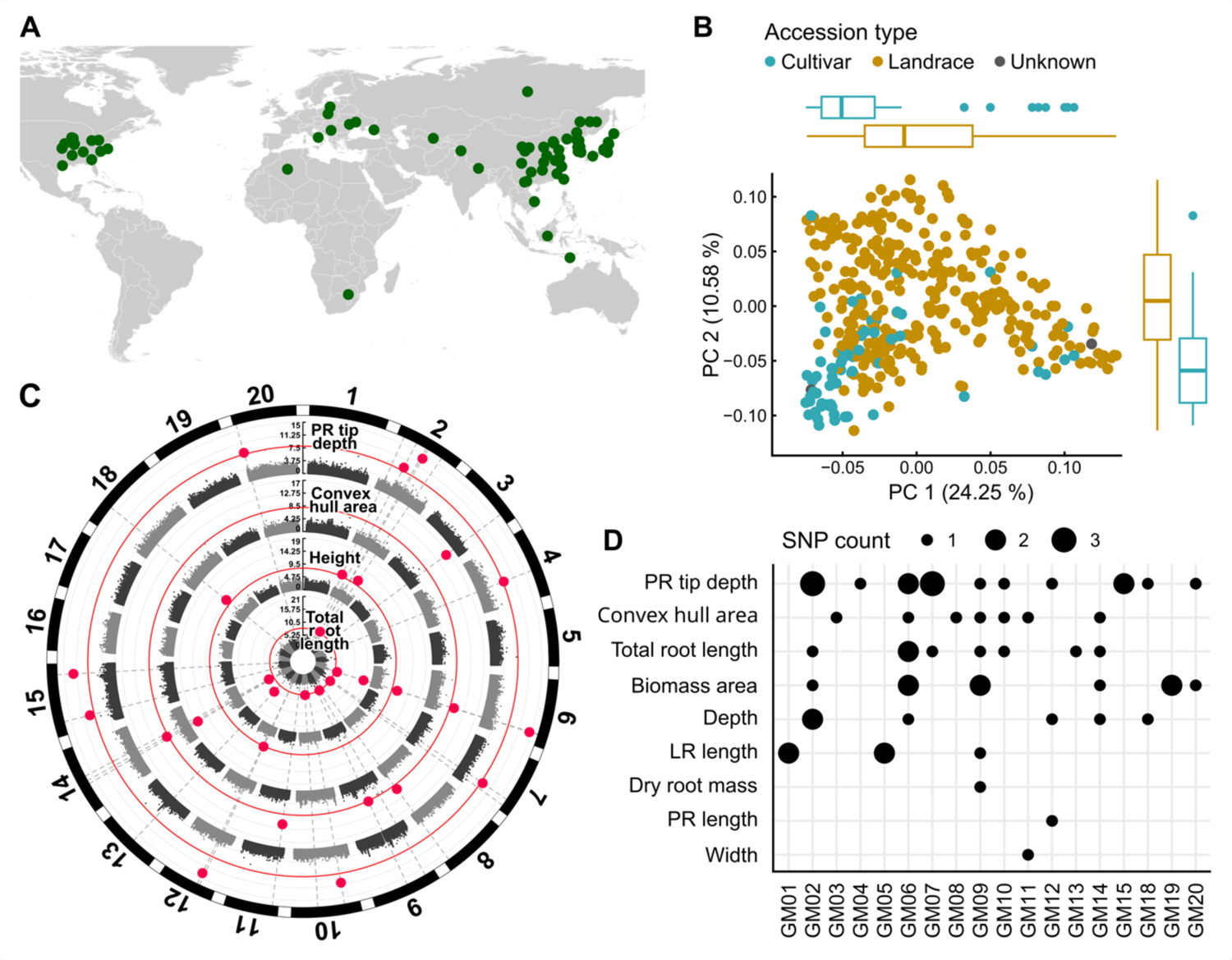
Genome-Wide Association Study (GWAS) identified specific and significant Single Nucleotide Polymorphisms (SNPs) associated with RSA across a panel of 371 soybean accessions. A) The locations of accessions used in this study originating from diverse regions across the world. Most of the samples are from Asia (∼60%). B) Principal Component Analysis (PCA) plot revealed population structure in soybean. C) Manhattan plot displaying significant SNPs as pink dots, using the FarmCPU model, associated with four traits from the inner to outer tracts: total root length, height, convex hull area, and primary root tip depth. D) Plot illustrating significant SNPs associated with specific traits, with larger dots indicating chromosomal regions housing a higher density of SNPs “hot spots”.

We made use of fastSTRUCTURE to explore the extent to which the RSA traits are influenced by underlying population structure and to perform a population structure analysis, which broadly correspond to the five countries of origin for these accessions (Supplementary Fig. 7a) (Raj, Stephens, and Pritchard 2014). We employed linkage disequilibrium (LD) analysis to determine the potential distance over which a SNP could be connected to a causal gene. We observed LD decay to a *r^2^* value of 0.2 at a distance of 300 kb (Supplementary Fig. 7b). This suggests long-range LD with the possibility of associated SNPs being linked with causal variants situated hundreds of kilobases away from the SNPs themselves.

A total of 30 model-RSA trait combinations were tested for statistical associations in our GWAS, which included 15 traits, each tested by Farm-CPU and BLINK models. In some cases, we observed overlaps or exact matches of associated loci (Figure 2c). In total, our analysis yielded 54 Bonferroni corrected (alpha = 0.05; p-value ≤ 1.03e-08) significant SNPs, of which 49 are unique loci distributed across 18 of the 20 soybean chromosomes (Figure 2d). QQ plots revealed no highly skewed p-values, suggesting that these results are reliable and not systematically biased by population structure (Supplementary Fig. 8). At each significantly associated SNP, we selected proximal genes within a 300 kb (LD distance), up to a maximum of 10 genes upstream or downstream, as potential candidates for functional prioritization. This resulted in a total of 633 GWAS candidates (GC) genes (Supplementary Table 1). We then incorporated relevant Gene Ontology (GO) terms associated with the GC genes and the description from the identified *Arabidopsis* ortholog. This approach aimed to provide insights into potential associated pathways (Supplementary Table 4).

Of the 49 unique significant SNPs, twelve fell within gene models: six in coding and five in non-coding DNA (Supplementary Table 2). Among these GC genes, seven have predicted *Arabidopsis* orthologs, while the function is unknown for the remaining genes. We examined their gene descriptions and revealed several genes with annotations related to regulating RSA. For example, Glyma.02G149100, which harbors a primary tip depth-associated SNP, Gm02:15770618, in the first coding region, is one of four soybean orthologs of the *Arabidopsis* glutathione peroxidase 4 (GPX4) (Passaia et al. 2014). Glyma.11G062900, one of eight soybean orthologs of *Arabidopsis* clathrin heavy chain 1 (CHC1), contains a SNP, Gm11:476025, in the 22nd coding region. The *Arabidopsis* mutant ortholog, *chc1*, demonstrated an increase in primary root length (Ormancey et al. 2020). Glyma.02G191500, one of two soybean orthologs of *Arabidopsis* DIHYDROFOLATE SYNTHETASE FOLYLPOLYGLUTAMATE SYNTHETASE (DHFS-FPGS) homolog C, contains a SNP, Gm02:38012571, within the 13th intron. Earlier research in *Arabidopsis* shows that maintaining folate levels in the leaves is required for maintaining plant metabolic homeostasis.

Consequently, the *Arabidopsis* ortholog, FOLYLPOLYGLUTAMATE SYNTHETASE 2 (*FPGS2*) has been associated with shorter root length (Zhang et al. 2023; Mehrshahi et al. 2010). Thirty-seven significant GWAS SNPs were found outside of gene models. Total root length and primary tip depth were associated with SNPs on seven different chromosomes (Chromosomes 2, 6, 7, 9, 10, 13, and 14 for total root length, and Chromosome 2, 5, 10, 12, 15, 18, and 20 for primary tip depth) (Supplementary Table 3). Conversely, certain traits such as dry root mass, primary root length, and width were solely associated with loci on a single chromosome (Figure 2c). Past studies have found RSA traits to be polygenic (LaRue et al. 2022), and these results are consistent with the suggestion that multiple GC genes distributed across the genome collectively influence traits, such as total root length and primary root tip depth, in RSA.

We also identified the presence of pleiotropic loci in SNPs that were associated with multiple traits. Specifically, the biomass area and primary root tip depth were associated with the same significant SNP loci (Gm06:17855981 and Gm07:35512631 respectively) in both the FarmCPU and Blink models. Additionally, two SNPs (Gm09:22884486, Gm06:41838269) were significant in multiple model-trait analysis, including FarmCPU-biomass area, FarmCPU-convex hull area, and FarmCPU-total root length. This may suggest a causal relationship behind the high correlations observed among these RSA traits (R^2^ = 0.9). Additionally, we identified a polymorphic gene, Glyma.06G250300. This gene encodes for a *ULTRAPETALA 1-LIKE* protein with unknown function. It was associated with two SNPs (Gm06:41838269 and Gm06:42199048) and exhibited correlations with FarmCPU-convex hull area and FarmCPU-total root length.

Furthermore, we identified “hotspots” where SNPs are in close proximity to each other at a particular locus such as: chromosomes 2 (37 Mb) and 6 (41 Mb) and are associated with depth, biomass area, convex hull, and total root length. These genomic loci are recognized based on having two or more significant SNPs and being within a 500 kb region (Supplementary Table 4). The observation that these SNPs are located in non-coding regions, with some concentrated together, implies the need for additional approaches to explore their impact on the genetic elements that govern the regulation of nearby genes.

### Differential expression and co-expression network analysis uncovered root-enriched network modules

By exploring natural variation in RSA through GWAS, we identified 633 promising gene candidates in close proximity to associated SNPs. 487 genes were predicted using Arabidopsis gene descriptions described to affect root morphology and function. The root traits measured in this study are quantitative traits expected to be regulated by multiple loci. In light of this, our aim was to investigate the connection between these genes by examining how they interact with each other through co-expression among the gene candidates. Employing weighted gene co-expression network analysis (WGCNA), we scrutinized gene expression patterns across diverse soybean datasets, summarizing expression landscapes across various organs and tissues.

Beyond elucidating potential functional interactions among the gene candidates, our method uncovered additional genes within the same coexpression network as the GWAS candidate genes. These genes in the shared network may play pivotal roles as regulators of RSA traits in soybean providing alternative targets for breeding or engineering. Our analysis first focused on compiling 71 soybean bulk RNA-Seq datasets accessed from the National Center for Biotechnology Information (NCBI) Short Read Archive (SRA) (Supplementary Table 5). This included 27 root, 18 leaf, 3 hypocotyl, and 3 shoot meristem datasets. Only data specifically labeled as the Williams82 variety were used in this analysis, and the data were aligned to the Williams82 reference genome (v4.0) (Figure 3a, Supplementary Fig. 10). We performed differential expression analysis across datasets using DESeq2 to identify differentially expressed genes (DEGs) enriched in specific tissues, with a significance threshold set at a p-value of <0.05 (Figure 3b, Supplementary Fig.10).

**Figure 3.**
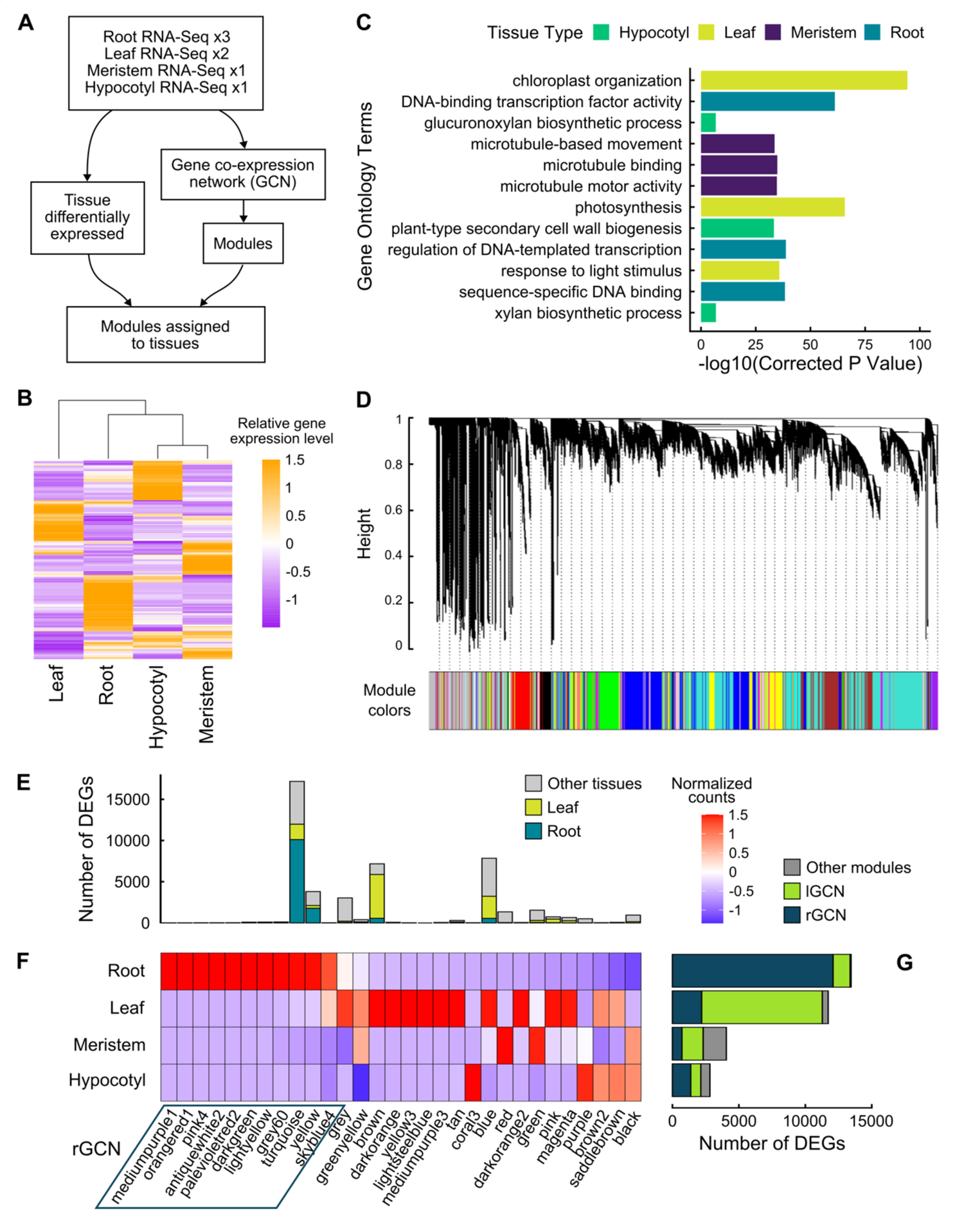
Differential gene expression and Weighted Gene Co-expression Network Analysis (WGCNA) identified root gene co-expression modules. A. Schematic of RNA-seq datasets and analysis. B. Heatmap of relative gene expression levels across tissues (10% most variable genes). C. Top Gene Ontology (GO) terms for Differentially Expressed Genes (DEGs) across four distinct tissue types. D. Identification of co-expressed modules using Weighted Correlation Network Analysis (WGCNA). Different colors represent multiple co-expressed gene modules of varying sizes. Identification of gene co-expression modules *via* hierarchical average linkage clustering. The color row underneath the dendrogram shows the module assignment determined by the dynamic tree cut. E-G. Enrichment of WGCNA modules with DEGs. (E) Total genes and root or leaf DEGs in each module. (F) Tissue enrichment of each module. (G) Total number of DEGs for each tissue.

We employed Gene Ontology (GO) enrichment analysis to test whether our analysis successfully led to the identification of gene sets that were enriched for tissue specific genes. Consistent with this hypothesis, patterns of enriched GO categories across different tissues were detected. For instance, the set of 10,670 leaf enriched genes was enriched for the GO categories chloroplast organization (corrected p-value 4.82e-95), photosynthesis (corrected p-value 1.94e-66), response to light stimulus (corrected p-value 1.57e-36), thylakoid membrane organization (corrected p-value 4.42e-34), and chlorophyll-binding (corrected p-value 1.24e-30). In the shoot meristem gene set of 2,721 genes, enriched GO categories included microtubule binding (corrected p-value 1.35e-35), microtubule motor activity (corrected p-value 2.16e-35), microtubule-based movement (corrected p-value 2.41e-34), and cell division (corrected p-value 1.40e-17). The hypocotyl gene set contained 1,856 genes and included enriched GO categories plant-type secondary cell wall biogenesis (corrected p-value 4.89e-34), xylan biosynthetic process (corrected p-value 1.52e-07), and glucuronoxylan biosynthetic process (corrected p-value 1.79e-07) (Figure 3c). Taken together, these results suggest that our analysis successfully identified tissue-enriched gene sets, as evidenced by the alignment of GO terms with the respective associated tissues.

We constructed a gene co-expression network (GCN) to identify putative network modules using WGCNA. We created a cluster analysis diagram to visually separate the samples and determined the optimal threshold for our analysis (Supplementary Fig. 11). The network construction resulted in a total of 99 distinct modules, each assigned to different colors (Figure 3d). Within these modules, we identified 30 containing DEGs (Figure 3e-g). We counted the DEGs of each tissue in each GCN module to identify the modules most relevant to root genes, sorting the modules based on the tissue most predominant in each module (Figure 3e-g, Supplementary Fig. 12). Among these modules, 11 were enriched for root genes, with “turquoise and “yellow” being the largest. The turquoise module contained 10,116 root DEG of 17,175 total genes (59%), while the yellow module had 1,779 root DEG of 3,805 total genes (47%). The turquoise module exhibited a close association with regulatory components, evident from the GO terms such as metal ion binding (corrected p-value 3.14e-50), DNA-binding transcription factor activity (corrected p-value 1.18e-39), and protein serine/threonine kinase activity (corrected p-value 1.40e-27). Additionally, the yellow module was connected with defense networks, indicated by GO terms such as response to chitin (corrected p-value 1.88e-62), cellular response to hypoxia (corrected p-value 2.98e-62), and response to wounding (corrected p-value 2.09e-22). We next adapted the PyGNA geneset network analysis software to further analyze the GCN (Fanfani, Cassano, and Stracquadanio 2020) (Supplementary Fig. 13). We extracted and summarized a subnetwork of the 11 root modules, termed the Root Gene Coexpression Network (rGCN) (Supplementary Fig. 14). The component size and degree distributions of the rGCN were similar to those of the full GCN.

### Network analysis prioritized GWAS candidates and specific root-enriched network modules for subsequent investigation

In our analysis, we refined the initial 633 GWAS candidate genes by focusing on only those differentially expressed in root compared to those differentially expressed in leaf, hypocotyl, shoot meristem or across several tissue types (Figure 4a). This subset of differentially expressed GWAS candidates (DE-GC) was distributed across 13 of the 99 WGCNA modules (Figure 4b-d). We identified 142 root differentially expressed GWAS candidate genes, which we termed rDE-GC, across 40 SNPs (Supplementary Table 6). Similarly to the overall set of rGCN (Figure 3f), the rDE-GC were prominently concentrated in the turquoise and yellow modules (Figure 4c).

**Figure 4.**
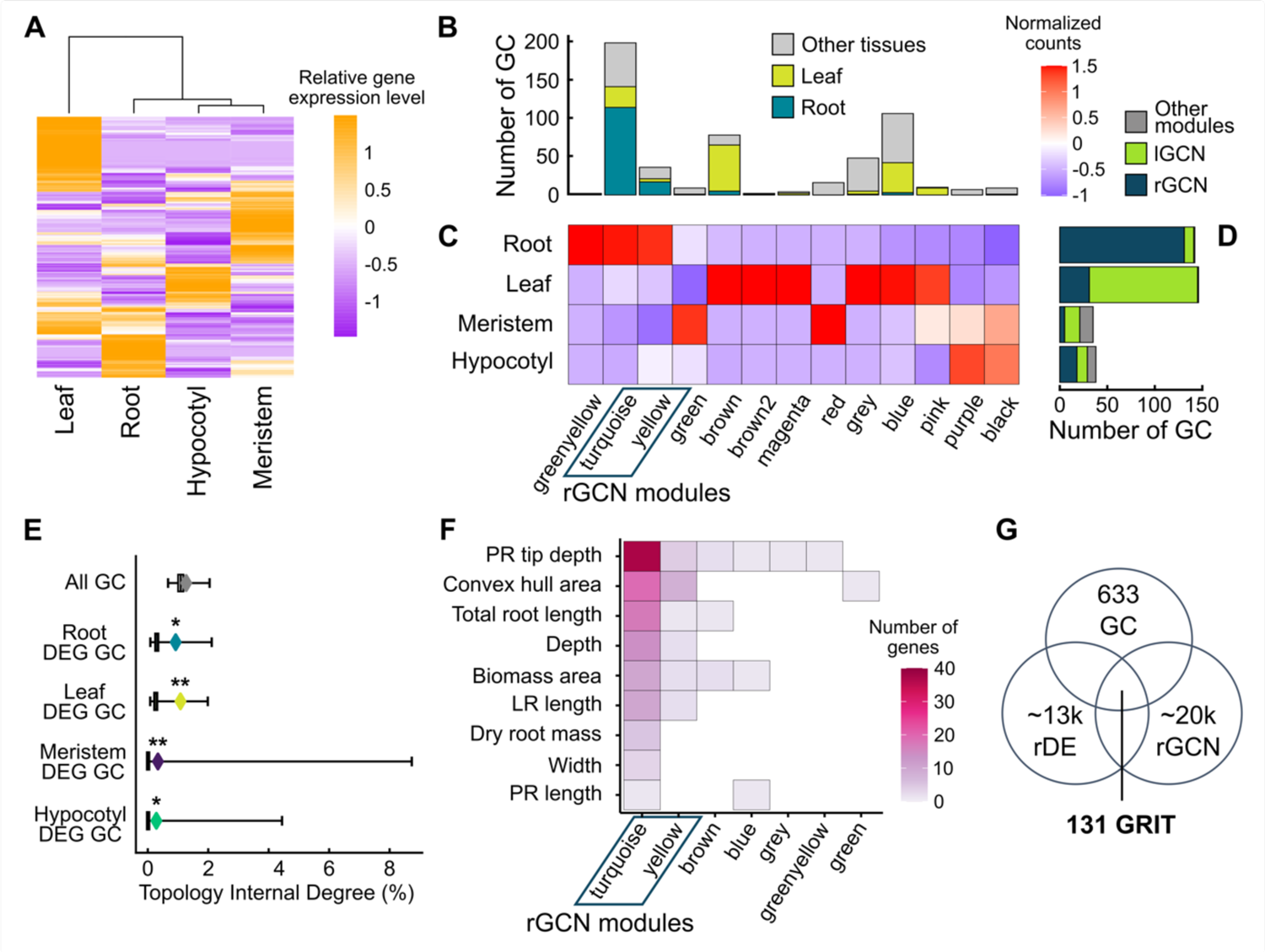
A high number of differentially expressed GWAS candidates (DE-GC) are concentrated in the turquoise subnetwork. A. Heatmap of relative gene expression level across tissues (GWAS candidates). B-D. Module enrichment of GWAS candidates. (B) Total GC and root or leaf DE-GC in each module. (C) Tissue enrichment of each module. (D) Total number of DE-GC for each tissue, with membership in root or leaf GCN modules shown. (E) Network topology internal degree test (%) tested for potential subnetworks between genesets and co-expression modules identified using WGCNA. “All GC” encompasses all GWAS candidates. DE-GC indicates differentially expressed GWAS candidates from root, shoot meristem, leaf or hypocotyl tissues. A significant association (P<…) is indicated by (*). (F) Heatmap of per-trait GC membership across GCN modules. Color bar indicates the number of genes. (G) Venn diagram illustrating the process of identifying genes associated with traits (GRIT) genes. The filtering process involves selecting GC genes, root genes that are DEGs for root vs. other tissues and present in a root-enriched gene coexpression network (rGCN).

Next, we explored the significance of rDE-GC and other DE-GC enrichment in modules by investigating their connection to network topology. Our goal was to determine whether these genes collaboratively functioned within specific modules, and if any of these modules exhibited tissue specificity. We employed the Topology Internal Degree (TID) test of PyGNA to quantify interconnectedness of gene sets, to assess whether the average internal degree of a gene set was greater than expected by chance. The internal degree statistic represents the average number of edges shared by genes within the set, indicating their connectivity within the network. Upon analyzing all GC genes collectively, we observed that the set was not topologically significant (p-value=0.07), likely due to the distribution of GC genes across multiple modules within the GCN. However, when we focused on the expression of tissue-specific DE-GC in the root, leaf, hypocotyl, and shoot meristem, the results were topologically significant (p-value<0.008). This indicated that the subsetted genes formed cohesive clusters within distinct modules, indicating their connectivity to specific modules within the GCN (Figure 4e). Our findings demonstrated the feasibility of identifying specific tissue-associated modules within the broader context of the GCN by discerning the connections of DE-GC within these modules.

Additionally, we delved into the rDE-GC set across 9 traits. Notably, the turquoise and yellow modules of the rGCN stood out as the most predominant, containing GC associated with nine and six traits respectively. Our observation suggested a higher level of complexity and interconnectedness within these modules (Figure 4f). Conversely, non-rGCN modules contained a minimal number of rDE-GC.

Building on our findings, we implemented an improved filtering strategy aimed at narrowing down our gene candidates. This approach was designed to minimize potential false positives and negatives from our GWAS analysis. The specific steps of this filtering process are visually depicted in a Venn diagram, illustrating the prioritization of GC genes (Figure 4g). This refined filtering involved narrowing down the pool of root differentially expressed GWAS candidate genes (rDE-GC) to those specifically present in a gene coexpression network enriched in root-related DEGs (rGCN) (Supplementary Table 7). Through this filtering process, we identified 131 genes meeting these criteria and referred to them as Genes Related to Identified Traits (GRIT).

### Exploring cell specificity of RSA trait regulation through single nuclei sequencing

Single-nuclei and single cell RNA sequencing (snRNA-seq/scRNA-seq) in plants has become a powerful technique to identify specific cell-types in complex tissues (D. Zheng et al. 2023; Bawa et al. 2022). Gene expression data at a high resolution can provide valuable insights into gene function and also be used to guide decisions when aiming to develop future crops with precise alterations in gene activity. We performed snRNA-seq on six-day-old whole root material of Williams82 soybean seedlings to gain further insight into the candidate genes identified in this study and to deepen our understanding of the soybean root expression landscape. After filtering for low quality nuclei, we obtained a dataset comprising gene expression data across 17,636 high quality nuclei capturing the expression of 47,095 transcripts. Following dimensional reduction and uMAP clustering, we identified 10 main clusters of nuclei with significantly unique transcriptional landscapes (Figure 5a). Due to the absence of specific markers for the majority of cell types in soybean, we employed a dual-strategy approach to annotate cell clusters within the Soybean Cell Atlas. Initially, a subset of clusters was classified utilizing established marker genes derived from recent studies (Liu et al. 2023), these clusters were annotated with the most likely cell-type identities compared with Arabidopsis (Supplementary Fig. 15). This annotation delineated several key cellular structures, including the metaphloem (cluster 12), cortex (clusters 0, 3, 6, and 9), epidermis (clusters 7 and 8), vascular bundle (cluster 11), and the xylem pole pericycle (cluster 2). Subsequently, we leveraged orthologs of well-characterized Arabidopsis marker genes to annotate the remaining cell clusters, which comprised the pericycle (cluster 1), xylem (cluster 10), and endodermis (cluster 5) (Supplementary Fig. 16). We generated cell-type specific gene sets by ranking genes according to variance in average gene expression across cell-types, selecting the top 10% most variable genes, and assigning each to 1 or 2 cell-type groups where their normalized expression value exceeded a constant threshold (Figure 5b, methods). The gene sets included 465 cortex-specific genes, 768 pericycle-specific genes, 410 vascular bundle and xylem pole pericycle-specific genes, 427 stele-specific genes, 672 endodermis-specific genes, 751 epidermis-specific genes, 886 xylem-specific genes, 840 vascular bundle-specific genes, 759 metaphloem specific genes, and 722 cluster13 specific genes (Supplementary Table 8).

**Figure 5.**
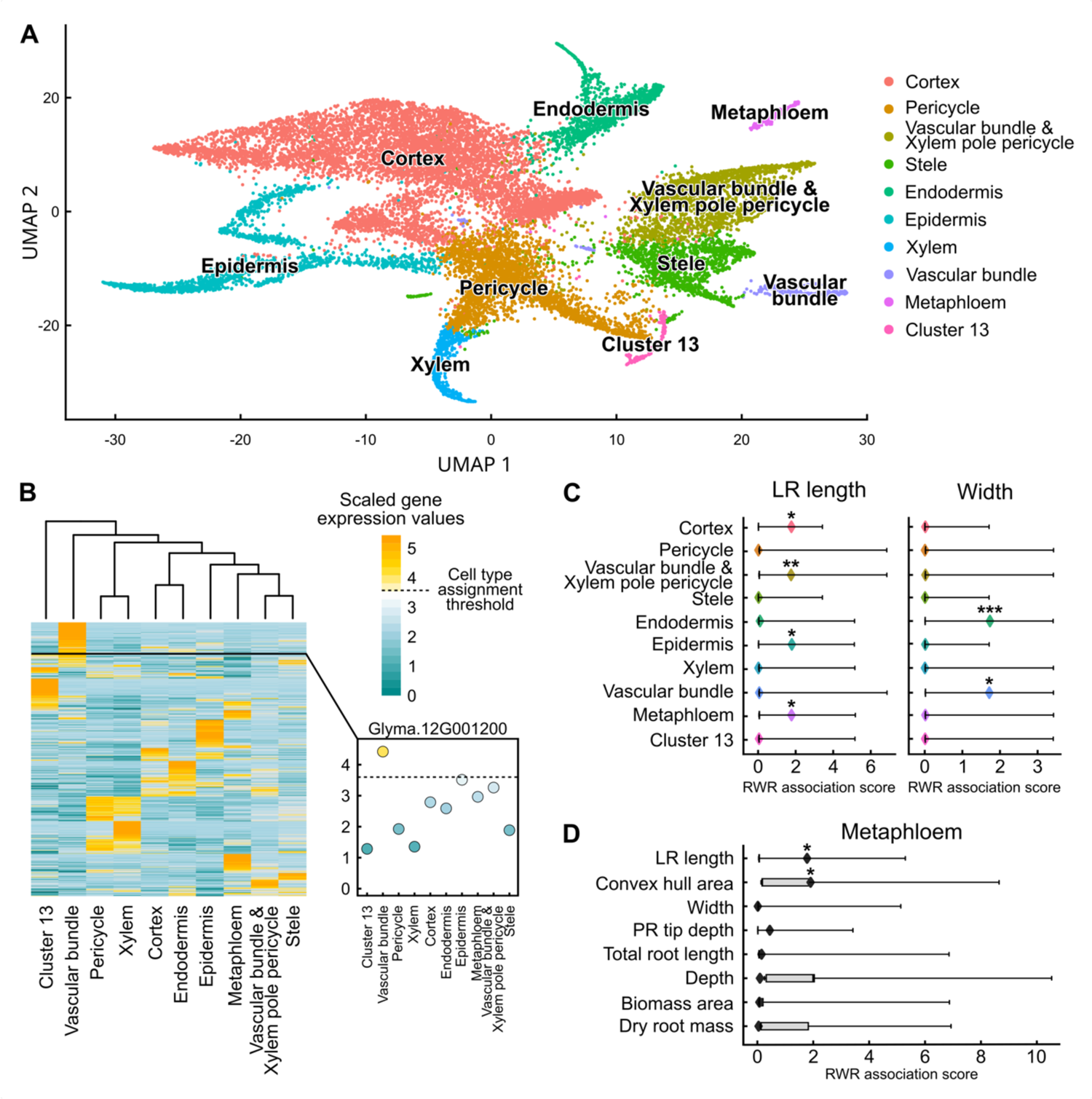
Single nuclei RNA-sequencing (snRNA-seq) enhances the resolution for spatial expression patterns of GWAS gene candidates, revealing insights into their cellular localization and potential functions. A. Uniform Manifold Approximation and Projection (UMAP) visualization illustrating distinct single-cell clusters, each representing a different cell-type. B. Heatmap depicting the expression levels of the top 10% most variable genes identified from a single-cell experiment, highlighting genes specific to different cell-types. Scaled expression values are shown. C. Boxplot of Random Walk with Restart (RWR) association test from PyGNA comparing GRIT genes separated by GWAS trait to the genes expressed from snRNA enriched in the metaphloem cell-type. D. Boxplot of RWR association test comparing GRIT genes for Lateral root length and Width traits to cell-type genesets. For both C and D, Boxes show null distributions, with whiskers showing the full range. Diamonds indicate observed values. * = p-value <0.01, ** = p-value <0.005, *** = p-value <0.001.

We found that of the 131 GRIT genes, 117 of them were expressed in the snRNA-seq dataset, so we restricted subsequent analysis of GC to these 117 (Supplementary Table 6). We identified several notable genes within these gene sets with previously identified root-related functions based on gene descriptions in the putative *Arabidopsis* ortholog. For example, Glyma.19G076800, is orthologous to AT5G40780 in *Arabidopsis* where it codes for LYSINE HISTIDINE TRANSPORTER 1. Our findings show that this specific gene is a candidate for biomass and is exclusively expressed in vascular bundles. Glyma.06G249700, an ortholog of AT1G54890, is a candidate for total root length, is expressed in all cell-types, and is related to Late Embryogenesis Abundant (LEA) proteins. Additionally, Glyma.06G230300, an ortholog of AT1G16890, is a candidate for primary tip depth and expressed in all cell-types, and encodes UBIQUITIN CONJUGATING ENZYME (UBC36/UBC13B), a protein involved in root developmental responses to iron deficiency in *Arabidopsis* (W. Li and Schmidt 2010). Taken together, these data point to our ability to identify genes related to roots within our single-cell data and that our findings are consistent with our current filtering approach.

We next proceeded to test for associations between specific cell-types and the networks where GRIT genes might be located. Our objective was to understand the potential co-expression patterns within the identified cell-types. We employed the Random Walk with Restart test from PyGNA to test for topological association of the GRIT gene sets and the cell-type gene sets within the context of the rGCN. Our analysis revealed several notable associations across different traits and cell-types (Supplementary Fig. 17). Of the different associations, we found that lateral root length exhibited the highest number of associations across multiple cell-types. Furthermore, the root width trait demonstrated a highly significant association with the endodermis (Figure 5c, Supplementary Fig. 18). This finding suggested that multiple cell-types may govern lateral root development while the endodermis may play a dominant role in governing root width. Variations in lateral root length and root width indicate that root traits might be regulated by multiple or a single cell-specific subnetwork and our analysis methods may potentially be instrumental in elucidating the cell-type specific genes governing these traits.

Additionally, our findings showed an exclusive association between the root metaphloem cell-type and lateral root length (Figure 5d). We determined that the metaphloem cell-type specific GRN was the only cell-type specific GRN associated with lateral root length with no significant association with other traits (Figure 5c). This suggested a potential metaphloem-specific role in influencing RSA through lateral root length. Our findings emphasized that our approach is useful in unraveling the complex regulatory pathways that govern RSA (Supplementary Fig. 19). The finer resolution provided by snRNA-Seq allowed us to identify potential cell-type-specific regulation in RSA in soybean. Analyzing genes within cell-specific subnetworks provides valuable insights into the molecular mechanisms governing variations in root width and lateral root length variation in soybean.

### Investigations into cell-specific networks identified gene networks associated with RSA control in the endodermis and metaphloem cell-types

Having identified subnetworks associated with specific and distinct sets of cell-types, our investigation aimed to determine whether the identified genes exhibited gene expression activities that could provide insights into their functions. Our initial focus was on the endodermis-root width associated subnetwork, which is a component of the larger turquoise network, and held the highest significance in our analysis. The endodermis surrounds vascular tissues and creates diffusion barriers to regulate the movement of water-soluble ions, protecting the root from the external environment. Within the endodermis subnetwork, we pinpointed two GRIT genes associated with root width, specifically linked to the SNP on Chromosome 11:31034808.

Glyma.11G176302 is a predicted ortholog of AT3G28050, a *USUALLY MULTIPLE ACIDS MOVE IN AND OUT TRANSPORTERS 41/EamA-like transporter* and Glyma.11G176301 corresponds to AT2G26730, a *LEUCINE-RICH REPEAT PROTEIN KINASE.* We assessed subnetwork connections, describing the number of edges extending from a node to other nodes (Figure 6a, Table 3). This calculation identified several co-expression connections for the GRIT genes in the context of the rGCN. *GmUMAMIT41* displayed the highest degree of connectivity, registering a degree of 4, whereas *Glyma.11G176301* exhibited a degree of 3. Co-expressed are other GRIT genes: Glyma.09G099500 (the predicted ortholog of *Arabidopsis METAL-TOLERANCE PROTEIN 10 [MPT10]* with cation efflux activity, AT1G16310) and a degree of 2; Glyma.20G058500 (the predicted ortholog of *Arabidopsis WSS1/SPRTN TYPE REPAIR PROTEASE B [WSS1B*], AT5G35690); and Glyma.15G127200 (the predicted ortholog of *Arabidopsis NONEXPRESSER OF PR GENES 3*, AT5G45110), with a degree of 3. We found that *GmUMAMIT41* and *GmMPT10* are transporters that exhibit the highest expression in the endodermis cell-type (Figure 6b). This suggests a potential link between the genes in this subnetwork and their influence on root width regulation through the processes located in the endodermis.

**Figure 6.**
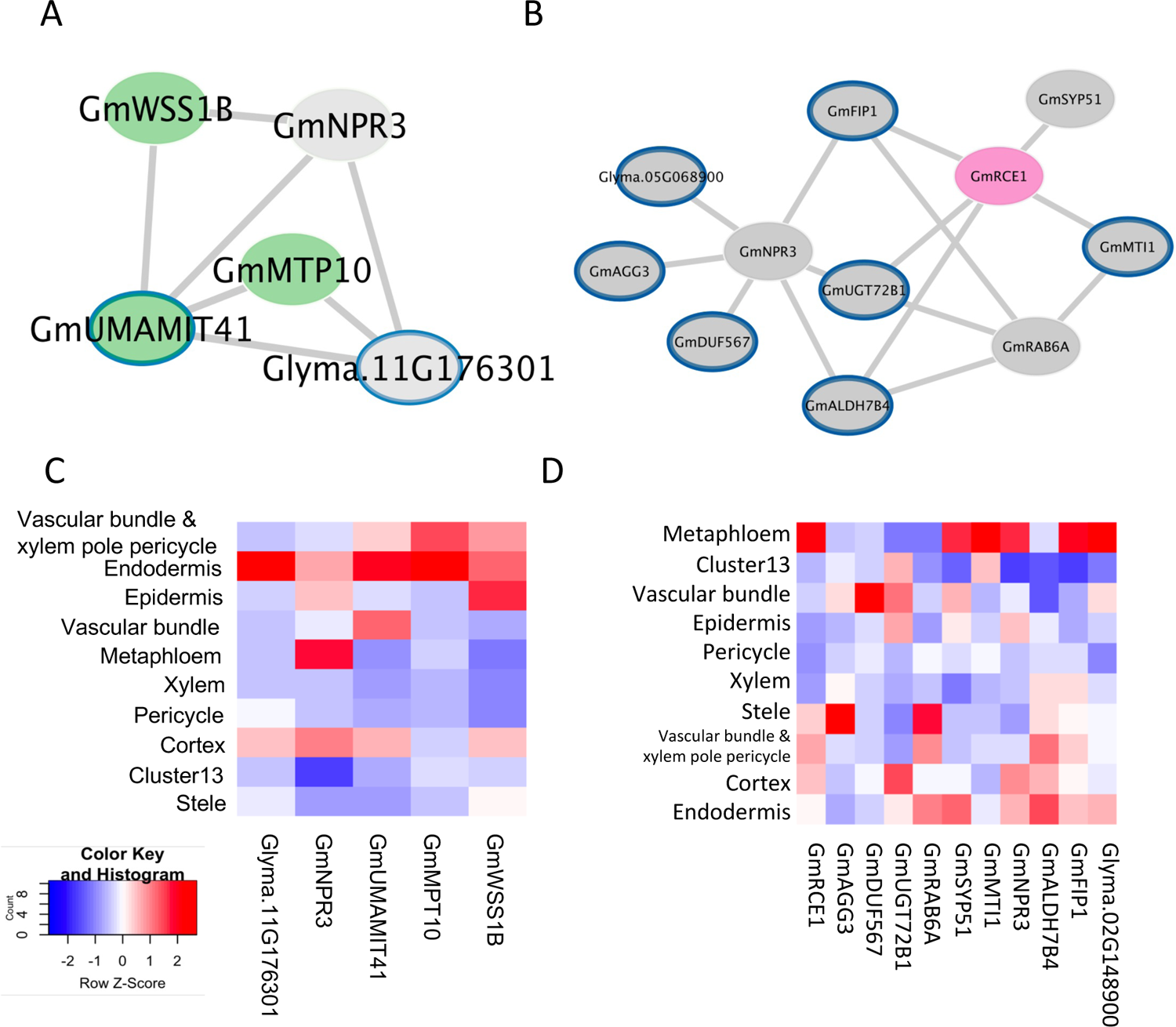
Identification of key gene co-expression network hubs for root width and lateral root development. A. Subnetwork highlighting GRIT genes (grey) associated with endodermis-specific GRIT genes in soybean (green) B. Subnetwork highlighting GRIT genes (grey) associated with metaphloem-specific GRIT genes in soybean (magenta) C-D. Heatmap depicting the average expression levels per cell for single-cell expression candidates.

Another notable subnetwork is the metaphloem network, which revealed a significant association between the metaphloem cell-type and GRIT genes related to lateral root length. The metaphloem plays a crucial role in transporting sugars and other solutes toward the root meristem (Graeff and Hardtke 2021). This subnetwork encompasses several genes co-expressed with genes in the metaphloem, including seven genes associated with lateral root length, two genes associated with convex hull, and two genes associated with primary tip depth (Figure 6c, Table 4). Among these genes, we identified Glyma.11G064000 (the predicted ortholog of *Arabidopsis RUB1 CONJUGATING ENZYME 1* [*RCE1*], AT4G36800), exhibiting a degree of 5. *RCE1* is involved in auxin signaling and the mutant exhibits morphological defects similar to those in mutants resistant to auxin (Dharmasiri et al. 2003). *GmRCE1* is connected to four other genes related to lateral root length: Glyma.09G070700 (the predicted ortholog of *Arabidopsis 5-METHYLTHIORIBOSE-1-PHOSPHATE ISOMERASE* [*MTI1*], AT2G05830) with a degree of 2; Glyma.01G074600 (the predicted ortholog of *Arabidopsis UDP-GLUCOSE-DEPENDENT-GLYCOSYLTRANSFERASE 72 B*1 [*UGT72B1*], AT4G01070), with a degree of 3; Glyma.08G322600 (the predicted ortholog of *Arabidopsis SYNTAXIN OF PLANTS 51* [*SYP51*], AT1G16240) with a degree connection of 1; and Glyma.09G069700 (the predicted ortholog of *Arabidopsis FH INTERACTING PROTEIN 1* [*FIP1*], AT2G06005) with a degree of 3.

Interestingly, among the genes we identified, those not directly related to lateral root length or metaphloem had the highest number of degree connections in this network. Glyma.15G127200, with the highest degree of 6, is a predicted ortholog of *Arabidopsis NPR3* (a paralog of *NONEXPRESSER OF PR GENES 1*, AT5G45110). It is connected to three genes related to endodermis and five other genes related to lateral root length. We discovered that numerous genes showed expression not only in the metaphloem but also in other cell-types. Interestingly, some of the co-expressed genes were not expressed the highest in the metaphloem (Figure 6d). The metaphloem serves as a physiological structure enabling the movement of substances (Hardtke 2023). Genes expressed in different cell-types might contribute to the functions of the metaphloem by affecting lateral root length when they are transported through this physiological structure.

Given the pronounced associations of lateral root length with multiple cell-types, we explored further by extending the network analysis to encompass its significant associations with all celltypes in the root including metaphloem, vascular bundle and xylem pole pericycle, cortex, and epidermis (Supplementary Fig. 20). This expanded analysis enabled the prioritization of 26 GRIT genes across four cell-types, which exhibited significant associations with lateral root length as well as other traits, such as convex hull area, primary tip depth, biomass area, depth, and total root length (Supplementary Table 11). Notably, *GmNPR*, *GmMTP10*, Glyma.18G226500 (*RPP39*), and Glyma.14G160100 (*SWEET3*) emerged as the top-ranking genes, with degrees of 16, 11, 11, and 10, respectively. Our analysis revealed several corresponding subnetworks associated with notable connections, offering a cell-type level understanding of the intricate network underlying these significant relationships.

## Discussion

We developed a novel 3D phenotyping method that non-destructively assessed RSA in 371 soybean seedlings. This method streamlined the analysis of root traits, resulting in a more high-throughput dataset that has greater scale and complexity in measurements when compared to other published methods. We subsequently conducted a GWAS involving 15 different RSA traits, 12 directly computed and 3 compound traits. This resulted in 54 significant SNPs supported by at least one GWAS model across 9 traits. Our network analysis refined 633 putative GWAS candidates by considering whether these genes were also differentially expressed in the root (rDE) and co-expressed (rGCN), revealing a total of 131 GRIT genes, 117 of which were expressed in the snRNA-seq dataset. We identified genes such as GmUMAMIT41, GmRCE1, and GmNPR3, along with their associated networks, as potentially influencing the variation in root width and lateral root length via the endodermis and metaphloem cell types.

Previous phenotyping methods have been constrained in their ability to fully capture the complexity of root growth due to constraints in measuring roots with sufficient spatial and temporal resolution. In field settings, environmental conditions may vary and introduce challenges in phenotyping, often relying on destructive measurements. Other approaches, such as employing 2D systems, fall short in capturing the entirety of RSA by not encompassing the natural growth patterns of roots in 3D. In contrast, our method allows for continuous observation without constraining the inherent growth process, a practice that has only been done so far in rice roots, with limited scale and data collection (Iyer-Pascuzzi 2010, Clark 2011).

We highlight the potential of utilizing snRNA-seq data to functionally prioritize GWAS candidates by assigning GWAS variants to their appropriate target genes, rather than solely based on proximity to the gene. Our investigation identified associations between endodermis and root width (widest span of the root structure), as well as metaphloem and lateral root length.

Furthermore, we pinpointed cluster 13 as a unique and distinct cell-type exclusive to soybean. In future studies, considering the potential differences between *Arabidopsis* and soybean, efforts to build robust validation of soybean markers, rather than relying on putative *Arabidopsis* orthologs for determining cell-types may reveal previously undiscovered cell-types in soybean.

Our integrative analysis in soybean represents the first model in soybean where GWAS, network analysis, and snRNA-seq were integrated to not only identify candidate genes but also elucidate their roles within specific cell-types, thereby providing understanding of how they contribute to morphological effects on RSA. In our broad soybean population, high LD and large haplotype blocks make it difficult to pinpoint the causal gene responsible for a GWAS association (Supplementary Fig. 7) (M.-S. Kim et al. 2021; Hyten et al. 2007; Chandnani et al. 2023). Our approach aims to extract insights into the functional roles of potential causative genes to improve the association of SNPs with their respective causal genes. While GWAS prioritizes candidate genes linked to target traits, network analysis reveals the interconnectedness of co-expressed genes. Additionally, snRNA-Seq suggests physiological function of specific genes or gene sets according to their expression across cell-types, facilitating the precise identification of causal genes. Our findings highlight the significance of this integrated method, especially in the context of crops or plants marked by high LD, such as soybean.

In comparison to earlier GWAS studies, our analysis aligns with past sample sets, typically involving several hundred samples ranging from 137 to 397 varieties (Salim et al. 2021; Dhanapal et al. 2020; Seck, Torkamaneh, and Belzile 2020; Prince et al. 2019; S.-H. Kim et al. 2023; Chandnani et al. 2023; Falk et al. 2020). These studies measured between seven and thirteen traits and identified two to 70 candidate genes, spanning a wide range of annotations (Salim et al. 2021; Dhanapal et al. 2020; Seck, Torkamaneh, and Belzile 2020; Prince et al. 2019; S.-H. Kim et al. 2023; Chandnani et al. 2023; Falk et al. 2020). Our analysis identified overlaps with two specific loci adding to the credibility of the discovered associations. We found that Gm18:51895181 from our study and associated with primary tip depth, is located 22,650 bp from one of the candidate SNPs: Gm18:51917831, which was associated with total root volume (Chandnani et al. 2023). Additionally, we found that a total root length SNP in our study (Gm09:7681193) is near (∼11kb) two candidates for the trait “number of forks” (NF) in (S.-H. Kim et al. 2023) (Gm9:7691924, Gm9:7699360), suggesting a potential quantitative trait loci (QTL) region for NF and number of tips (NT). It’s plausible that the variations in our dataset are from differences due to environment and developmental stages considered during our measurements.

Our data revealed insights into the primary subset of cell-types and genes involved in RSA in soybean. Notably, the strongest association from our analysis connected root width and endodermis. Root width is a frequently overlooked trait as most studies focus on length and position. However, improving root width could hold potential for optimizing soybean planting by maximizing soil nutrient utilization between plants. We identified a subnetwork of GWAS candidates expressed in the endodermis cell-type. Among them are amino acid and ion transporters, which aligns with the role of the endodermis in regulating water and nutrient movement to and from the vascular system. These findings could be valuable in understanding how variation in endodermis function could impact the development of root width.

Additionally, we identified associations of multiple cell-types with lateral root length, including cortex, epidermis, xylem pole pericycle, and metaphloem. For the metaphloem cell-type, lateral root length was the sole trait for which genes proximal to GWAS loci were topologically associated. Despite existing knowledge on lateral root emergence and the role of auxin, the complexities surrounding the regulation of long-distance systemic signals pose challenges for molecular studies (Geng et al. 2023). Given that the phloem serves as a specialized transport facilitator, the contribution to variations in lateral root length through genes may not be exclusive to the metaphloem. In this context, we identified a gene expressed in the metaphloem cell-type involved in auxin signaling which could illuminate the unique functions of metaphloem during lateral root development.

Considering the networks more broadly, we observed that GmNPR3 exhibits the highest degree connections across all three networks related to lateral root length, width, and metaphloem. Previous investigations in *Arabidopsis* have demonstrated that its ortholog, AtNPR1, potentially modulates lateral root abundance by mediating the antagonistic interaction between auxin and salicylic acid (SA) during *Pseudomonas* invasion (Kong et al. 2020). This implies that the crosstalk among various hormones, such as SA and auxin, aids plants in evading pathogen attack by regulating lateral root growth under biotic stress. Therefore, GmNPR3 likely establishes links to genes associated with plant defense in addition to influencing root width and lateral root length. Future gene edits to NPR3 could prove beneficial for increasing lateral root length, particularly if done in a cell-specific manner or by targeting its downstream targets.

Understanding genes within the context of networks provides a more comprehensive overview of how traits may be regulated and thus provides more information that can be used to guide crop editing efforts, minimizing unintended effects. Our approach contributes to a greater understanding of the regulation of complex traits within cell-specific networks, providing a more comprehensive view of the genes with pleiotropic function. Future directions may involve exploiting expression quantitative trait loci (eQTL) and increasing the genetic diversity of our plant population. Ultimately, our findings could lead to developing more effective multiplexing CRISPR gene edit techniques or markers that aid in marker-assisted selection for developing new soybean varieties with enhanced root systems.

## Materials and Methods

### Data and code availability

Supplementary Data can be found here: https://salk-tm-pub.s3.us-west-2.amazonaws.com/Sun_etal_supplementary_data/Sun_etal_supplementary_data.zip. The code to analyze the WGCNA network and single-cell data can be found here: https://gitlab.com/salk-tm/soybean-root-gwas/. RADYCL Segmentation pipeline for image analysis can be found here: https://github.com/Salk-Harnessing-Plants-Initiative/SSRAPC-Soy-Segmentation-Root-Architecture-Phenotyping-for-Cylinder.git. PyGNA2 is available on PyPI (https://pypi.org/project/pygna2/) and GitLab (https://gitlab.com/salk-tm/pygna2).

### Germplasm collection

A diverse set of 371 soybean plant introductions (PIs) was selected from the USDA Soybean Germplasm Collection (Valliyodan et al. 2021). The collection represents a wide genetic diversity with genotypes collected from over 20 different countries and includes maturity groups (MG) ranging from II to V. The population is mainly/majority have individuals/genotypes from China (222), USA (52), Korea (36), Japan (23) and Russia (11).

### Genotyping

Sequencing data was generated using previously described methods (Valliyodan et al., 2020). In short, trimmed paired-end Illumina reads from each sample were aligned to the Williams 82 reference (Glycine max Wm82.a4.v1) using BWA (H. Li and Durbin 2009) and Picard tools (Wysoker, Tibbetts, and Fennell, n.d.). GATK (https://github.com/broadinstitute/gatk/) was then used to call variants. Next, VCF tools (Petr et al., n.d.) was then used to select biallelic SNPs that were present in at least 50% of samples with a minor allele frequency of no less than 5%, a minimum depth per sample (DP) of 10 reads, and minimum genotype quality score (GQ) of 10. This yielded 4,815,704 SNPs for downstream GWAS analyses.

### High throughput gel-based phenotyping for root system architecture

Soybean seeds were surface sterilized with chlorine gas (4mL HCL in 250 mL bleach solution) in a fume hood for 12 hours. One seed was sowed for each plastic cylinder filled with 170 ml of half strength Murashige and Skoog (MS) medium (pH 5.7) solidified using 0.8% phytagel. The dimensions of the plastic cylinders (Greiner Bio-One Polystyrene Container, Product No.: 968161) were 110 mm height and 68 mm diameter with volumetric capacity of 330 mL. Eight individual replicates were generated for each accession and placed in a randomized block design in a greenhouse with natural and artificial lighting (28 ± 8 °C, 14-h photoperiod). The images of the root systems were acquired 5- or 6-days post germination using an imaging system we termed Root Architecture 3D Imaging Cylinder (*RADICYL*).

The imaging system consisted of a telecentric lens mounted in a Basler acA2000-50gm GIgE camera, an aquarium, and a light source mounted on an optical breadboard. Gel-filled cylinders containing the soybean seedlings were placed in the water-filled aquarium that was placed on a turntable utilized to position the cylinders. This setup was back-illuminated by a near-infrared light source (Supplementary Fig. 1) and a rotational series of images of the cylinder were acquired (72 images in 5° steps). After image acquisition, seedlings were pulled out from the gel. Roots were separated from shoots and pinned on a cardboard with respective labels and then dried for 72 hours at 50°C. The dried roots were then weighed using a fine scale.

### Image analysis, skeletonizing and trait extraction

We employed UNet++, a deep learning model, to segment primary/lateral roots from images collected using *RADICYL*: the UNet++ architecture was optimized using Adam optimizer with an initial learning rate of 0.0001 for 60 epochs. We used ResNet101 as the encoder of the UNet++ model and softmax2d as the activation function of the last layer. The best validated model with the highest IoU score was saved as the trained model. During the segmentation phase, 72 images for each cylinder were cropped and used for primary/lateral roots segmentation based on the trained UNet++ framework. Roots on the images were then manually labeled using LabelMe (https://github.com/wkentaro/labelme) with three classes of labeled pixels: primary roots, lateral roots and background. The training-set contained 90 *RADICYL* captured images with a size of 2048 x 1080 pixels that were randomly selected and not part of our analysis. We cropped the images to a size of 990 x 860 pixels centered on the cylinder to remove pixels outside of the cylinder area.

To skeletonize the roots (primary roots and lateral roots) from segmented images, the function pcv.morphology.skeletonize in PlantCv was used. The python library, OpenCV, was used to measure convex hull; the Hough Line Transform function from OpenCV was used to detect line segments originating from a given base of a lateral root on the primary root and ending on the tips of a lateral root, the average vertical angle for these short straight lines for all lateral roots was computed and output as vertical angle. During the prediction phase, 72 images for each cylinder were cropped and used for primary/lateral roots segmentation based on the trained UNet++ framework. The resulting black and white ground truth images were generated and cross checked with the manually annotated images. In total, nine different RSA-related traits were measured from each 2D image. A quality control step was performed by producing collages with segmented images of each plant belonging to a single genotype. A user then identified poorly germinated or abnormally grown plants and excluded them from the final analysis. The traits mean and median data of the final set of images were used in the subsequent analysis. Each trait data set was tested for normality utilizing the Kolmogorov– Smirnov test and applied transformations to data that failed the normality test using the BestNormalize package in R.

### GWAS analysis pipeline

GAPIT3 (Jiabo Wang and Zhang 2021) was utilized to test for associations between the SNPs from 371 samples and the 15 root traits. Using the median values for each accession and trait, plus the two most statistically powerful models (BLINK, FarmCPU) lead to 22 model-trait combinations being tested (gitlab.com/salk-tm/snake_gapit_gwas). BLINK and FarmCPU are both multilocus models, which incorporate the top three principal components derived from all the markers as covariates to reduce false positives. Additionally, BLINK iteratively incorporates associated markers as covariates to control for relationships among individuals. The associated markers are first selected using linkage disequilibrium, then optimized for Bayesian information content, and finally reexamined across multiple tests to reduce false negatives. Results from each model-trait combination were then pooled and treated as independent tests. P-values of less than 5% after Bonferonni correction for multiple testing (p = 0.05 / 4.1 million SNPs) were considered as significant. We extended the region of interest up and downstream of the significant SNP by 10 genes on each side.

### Construction and analysis of root gene networks

Gene expression data representing diverse tissue types of soybean was obtained from Short Read Archive (SRA). Samples were filtered for only samples related to the Wm82 accession which was a total of 71 gene expression samples were selected for further analysis. The raw gene expression data was preprocessed and normalized using Salmon and transcript counts were quantified (Patro et al. 2017). DESeq2 was used to identify differentially expressed genes for all four tissue types (Love, Huber, and Anders 2014). The WGCNA package in R was applied to construct a gene co-expression network (Langfelder and Horvath 2008; Chang n.d.). Network construction was performed using the *blockwiseModules* function for the entire dataset to calculate a pair-wise correlation matrix and adjacency matrix for each set of genes. We calculated the topological overlap matrix (TOM) for all pairs of genes in the co-expression network to quantify the strength of interconnection between two genes before defining the modules within a network. Using the modules, 10 was set as the minimum number of genes in the module and the threshold of cutting height (Supplementary Data). The gene modules were identified using the dynamic cut tree method and the modules with high similarity were combined to obtain 99 modules. Root-enriched modules were defined as the modules that had the highest number of root DEGs within the module. Initially, WCGNA generated a network comprising 219 million edges. After filtering only the root-enriched modules, this number was reduced to 154 million edges. Then by applying a threshold for the correlation coefficient (≥ 0.1), the network was further refined to include 10 million edges. A summary of the network was generated using PyGNA to investigate properties of the root-enriched subnetwork, including number of nodes (genes), edges (co-expression relationships), and degree (number of edges) of each node. We identified the root-enriched network to consist of 18,191 nodes, and 10,326,288 edges, with individual nodes having a minimum degree of 1 and a max degree of 7,516.

### Nuclei extraction and snRNAseq library construction

Williams 82 soybean seeds were sterilized in the same manner as the cylinder experiment above and grown on filter paper soaked in ½ MS for germination in a Percival growth chamber at 28°C/18°C and 12 hr/12 hr day night conditions. 7 days post germination, root material was harvested in liquid nitrogen and stored at −80 until further processing. For nuclei extraction (using methods described by (Lee et al. 2023)), 75 roots were ground to a powder using a cooled pestle and mortar and homogenized in 20 ml Nuclei Extraction buffer (NEB) (20 mM MOPS (pH 7), 40 mM NaCl, 90 mM KCl, 2 mM EDTA, 0.5 mM EGTA, Supplemented with *0.5 % SUPER RNase inhibitor,* 0.5 mM Spermidine, 0.2 mM Spermine, 1:100 dilution Roche Complete Protease Inhibitors).

Samples were then sequentially filtered through a 70 μm and then 40 μm cell strainer and centrifuged for 5 minutes at 700 rcf (all centrifugation steps were performed at 4°C). The liquid phase was removed using an aspirator and the pellet was resuspended in NEB + 0.1% triton. Samples were incubated on ice for 15 minutes before being centrifuged at 700 rcf for 5 mins. This step was repeated for a total of three washes. Following the third wash, the pellet was resuspended in 4 ml NEB. A density gradient was used to separate the nuclei. For this 1 volume of diluent (120 mM Tris-Cl pH 8, 150 mM KCl, 30 mM MgCl2) was added to 5 volumes 60% Optiprep to make a 50% density buffer. This 50% stock was then used to make 45% and 15% solutions using a second dilution buffer (400 mM Sucrose, 25 mM KCl, 5 mM MgCl2, 10 mM Tris-Cl pH 8). The gradients were made in 15ml tubes with 1 ml of 15% solution and 2 ml of 45% solution. Two ml of each sample was added to the gradient and centrifuged at 1500 rcf without breaks. Following centrifugation nuclei can be seen as a layer at the 45% mark in the gradient. These were collected into a 15ml falcon tube and centrifuged at 1000 rcf for 5 mins.

The nuclei pellet was resuspended in 1 ml NEB and nuclei were sorted using Hoechst stain. The sorted nuclei were centrifuged at 700 rcf for 5 mins and resuspended in 50 ul 1xPBS to ensure compatibility with 10X Genomics library preparation. Libraries were made using the Chromium Next GEM Single Cell 3ʹ Reagent Kits v3.1 according to manufacturer’s instructions. cDNA and final library quality were assessed with a Bioanalyzer D1000 DNA Chip (Agilent). and libraries were sequenced using the (Illumina) NovaSeq SP 100 cycle kit.

### Raw snRNA-Seq data pre-processing

The initial analysis of the raw snRNA-seq dataset was conducted utilizing Cell Ranger (6.1.2) mkfastq (10X Genomics). This process involves alignment of reads and generation of gene-cell matrices. Both genome and GTF annotation files of *Glycine max* were procured from Gmax.Williams82. The reference was constructed by executing the ‘cellranger mkref’ command with ‘-genome, -fasta, and -genes’ arguments. The ‘cellranger count’ command was used with ‘-id, -transcriptome, -fastqs, -sample, and -force-cells’ arguments to generate counts of single-cell genes.

### Cell clustering reconstruction via nonlinear dimensionality reduction

We next aimed to determine if candidate genes associated with root-specific network modules were expressed in the same cell-type. To this end, we performed single nuclei RNA sequencing on six-day-old whole root samples from the reference cultivar Williams82. This approach was instrumental in investigating the co-expression of gene networks within specific cell-types, offering a more refined understanding of the spatial and functional organization of genes in the root. In our analysis of Soybean snRNA-Seq datasets, we initiated the process by normalizing the UMI counts for each gene. This was achieved by dividing the UMI counts by the total UMIs in each cell, scaling the result by 10,000, and then applying a logarithmic transformation. We rigorously filtered the cells, setting stringent criteria based on the percentage of mitochondrial transcripts (percent.mt), the number of detected genes (nFeature_RNA), and the total mRNA molecules in each cell (nCount_RNA). Our thresholds were tailored to soybean datasets: nFeature_RNA per cell had to be more than 300 but less than 2500, percent.mt was capped at 1%, and nCount_RNA had to be between 300 and 4000. Following quality control, we retained 17,636 high-quality nuclei capturing the expression of 47,095 genes. Next, we identified the 4,000 highly variable genes using the FindVariableFeatures function in capturing the broader variability in transcriptomes. Dimensionality reduction was then performed via PCA using the ‘RnPCA’ function. To address batch effects, we employed *Harmony*, ensuring accurate integration and analysis of our snRNA-Seq data. For unsupervised clustering, we utilized the ‘FindNeighbors’ function with the top 20 PCs and the ‘FindClusters’ function at a resolution setting of 0.3. These steps are crucial in revealing inherent grouping patterns within the data.

The resulting clusters were visualized using the UMAP method, facilitating an intuitive understanding of the data’s underlying structure. With known soybean marker genes, orthologs of marker genes in Arabidopsis, we successfully identified cortex, pericycle, vascular bundle/xylem pole pericycle, stele, endodermis, epidermis, xylem, vascular bundle, metaphloem.

### snRNA-Seq data post-processing

10 cell-types were annotated based on known marker genes and gene expression profiles. Cell clusters “inner cortex”, “outer cortex”, and “potential cortex” were merged into the cortex cluster to simplify the analysis. 47,094 genes were sorted by their variance across cell-types, and the top 10% (4,709) most variable genes were selected for downstream analysis. For each variable gene, we scaled expression values to a standard deviation of 1 across cell-types, and a common mean across genes. We divided the range of scaled expression values into 3 equal bins, and for each cell-type chose genes with scaled expression values in the top bin as the cell-type specific gene set.

### PyGNA 2 analysis

To facilitate our hypothesis tests and visualizations, we developed a Python package based on the API of PyGNA, which we called PyGNA2 (Fanfani, Cassano, and Stracquadanio 2020). PyGNA2 was employed to summarize the gene networks. We used the ‘pygna2 summary’ function to obtain key network statistics, providing an overview of network characteristics. We employed the ‘pygna2 test’ function to assess the relationship between the root-enriched gene network and the single-cell gene sets. This function tests the associations between two networks using the Topology Random Walk with Restart test and the Topology Internal Degree test (TID). We performed all our final analysis with 8000 permutations. We visualized the results using ‘pygna2 cytoscape’ with the –minimal option to visualize subnetworks including specific cell-type gene sets, GWAS candidate genes, and genes on shortest paths between them.

### Network filtering and cytoscape analysis

We identified the largest connected component within the significant cell-type vs. trait network and designated this as “cell-type and trait”. We retained only the nodes corresponding to genes associated with cell-type and trait within the larger network component to simplify the network further. Node selection was guided by the 117 GRIT genes identified from filtering by root DEGs within root modules in single nucleus data. Additionally, we determined the degrees of connectivity for each subnetwork and determined that these central connecting nodes to be the most significant.

## Acknowledgments

We thank Emily Shane and Charidimos Georgousakis for phenotyping. This work was supported through the Salk Harnessing Plants Initiative (HPI) with funding from the TED Audacious, Bezos Earth Fund, and Hess Corporation.

## Declaration of interest

W.B. and T.M. are co-founders of Cquesta, a company that works on crop root growth and carbon sequestration.

---

**Supplementary Fig. 1:**
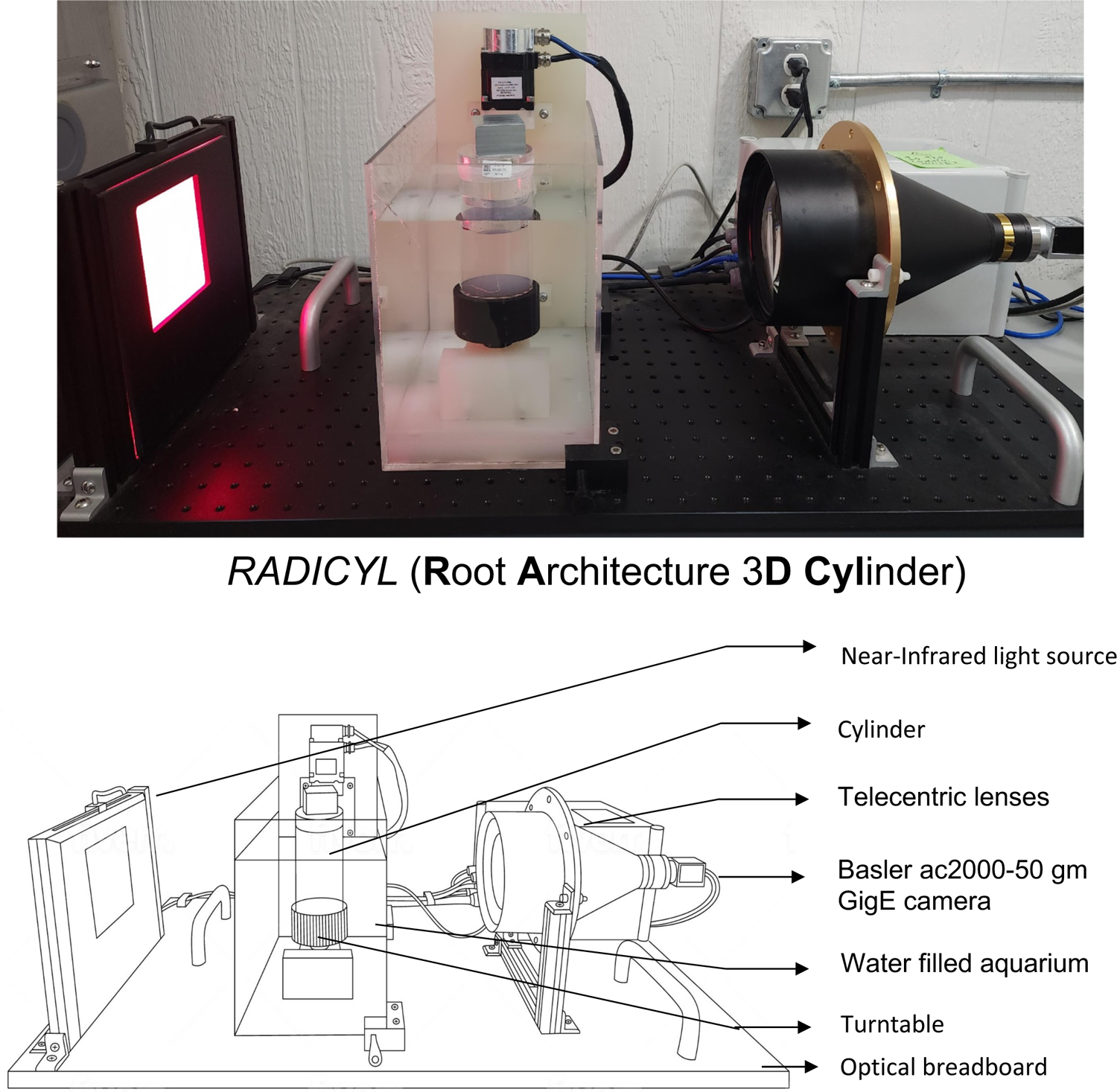
Root Architecture 3D Cylinder (RADICYL) - A High-Throughput Phenotyping System. The RADICYL system comprises key components, including an automated turntable equipped with a Basler acA2000-50gm GigE camera and a telecentric lens, positioned above an aquarium. Additionally, an optical breadboard-mounted light source is used to illuminate the subject. In the experimental setup, cylindrical samples are submerged in a water-filled aquarium, which is then placed on the turntable. A rotational series of images is captured, consisting of 72 images at 5-degree intervals. Subsequently, seedlings are carefully extracted from the hydrogel medium following image acquisition for dry biomass measurements.

**Supplementary Fig. 2:**
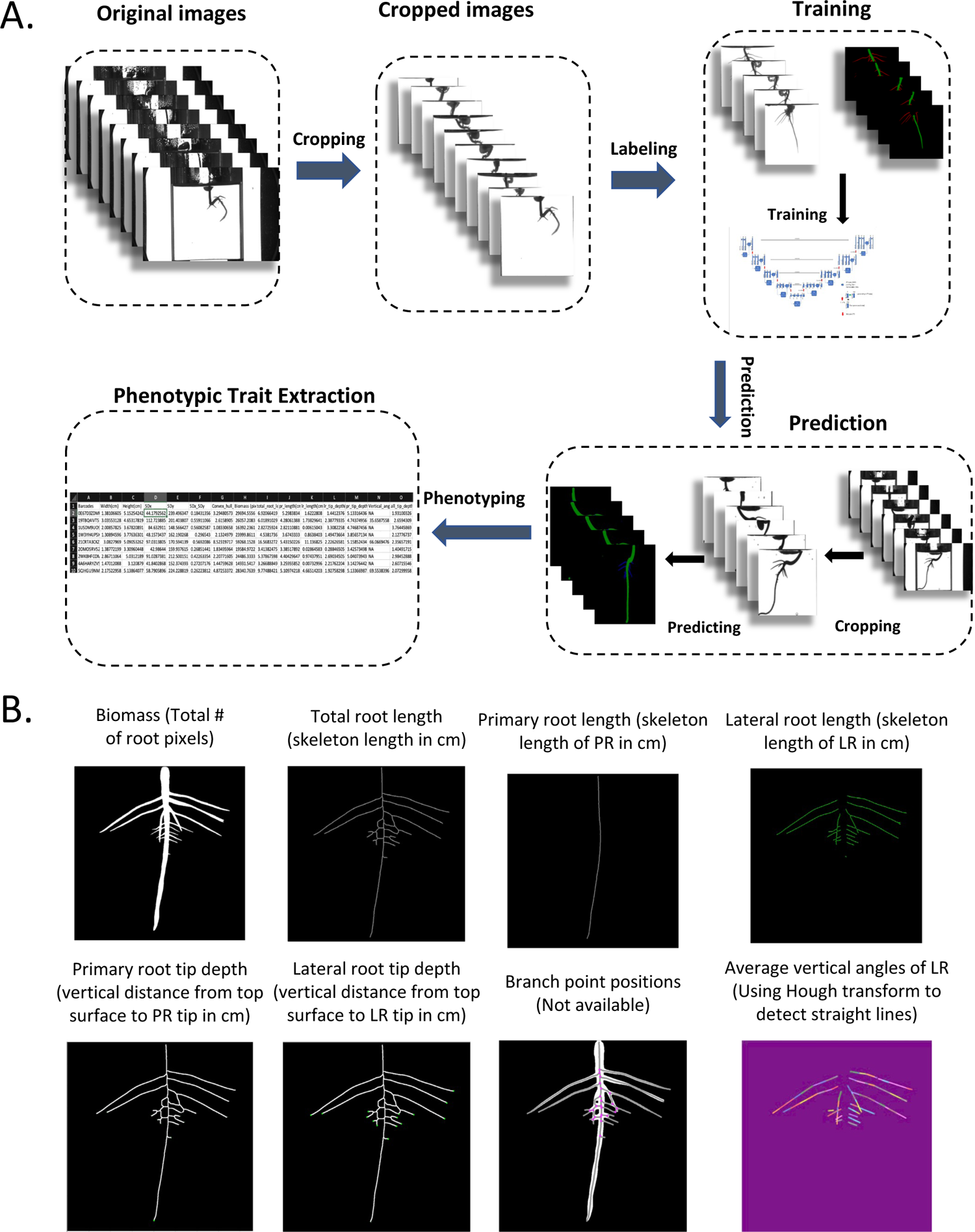
A. An illustrative example of the processing steps undertaken to extract the trait quantifications after capturing the initial images. B. Segmentation of primary roots and lateral roots was accomplished using the UNet++ architecture, a convolutional neural network (CNN) designed for semantic segmentation tasks. During the segmentation phase, 72 images from each cylinder were cropped and used for primary and lateral root segmentation based on the trained UNet++ framework.

**Supplementary Fig. 3:**
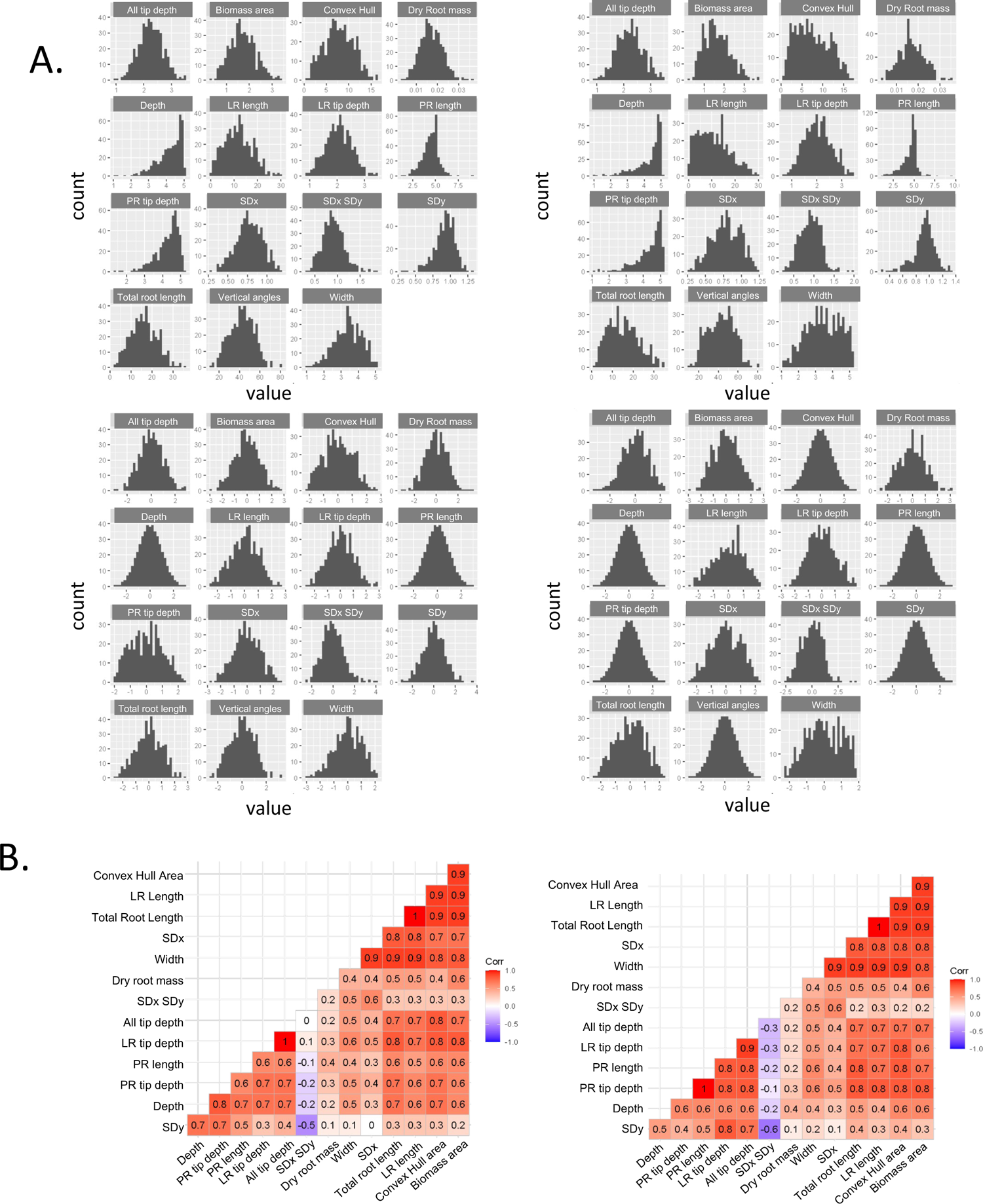
A. The histograms display the mean (on the left) and median value (on the right) for 15 RSA traits. To ensure the data met normal distribution criteria, adjustments were made to transform the data (displayed in the bottom left and right). For the GWAS analysis, the median values were used to avoid potential biases from extreme values that could skew the mean. B. In correlation plots, the left side illustrates correlations using the mean, while the right side does so with the median. Both plots emphasize strong connections between the various quantified traits, revealing a high level of correlation among them.

**Supplementary Fig. 4:**
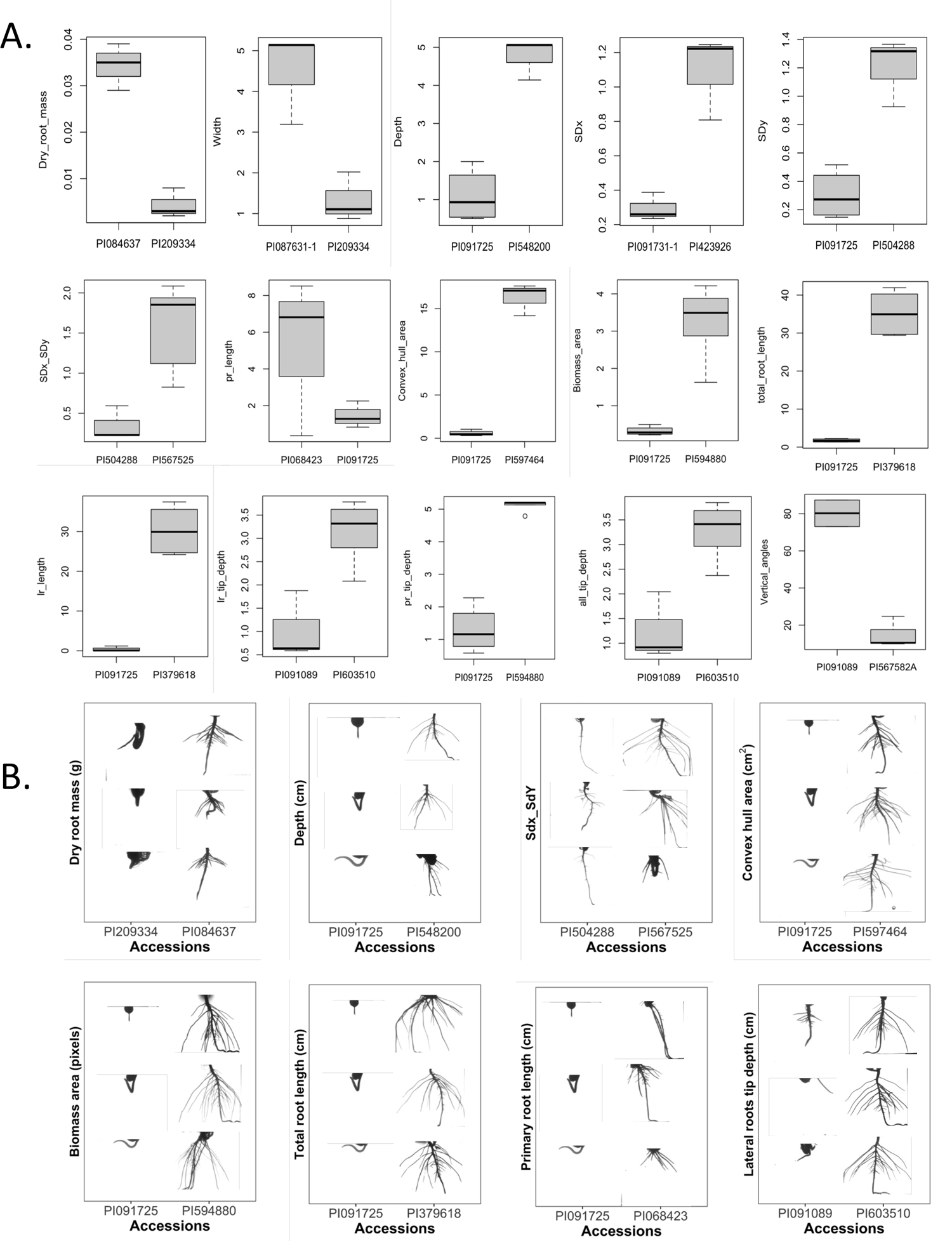
A. Box plots depict the two highest and lowest accessions associated with the median of each trait. Each box plot summarizes the replicates for the respective trait. Notably, several accessions exhibit both high and low associations across multiple traits, revealing complex relationships within the dataset. B. Phenotypic differences in two extreme, the highest and lowest accessions associated with the median of few traits.

**Supplementary Fig. 5:**
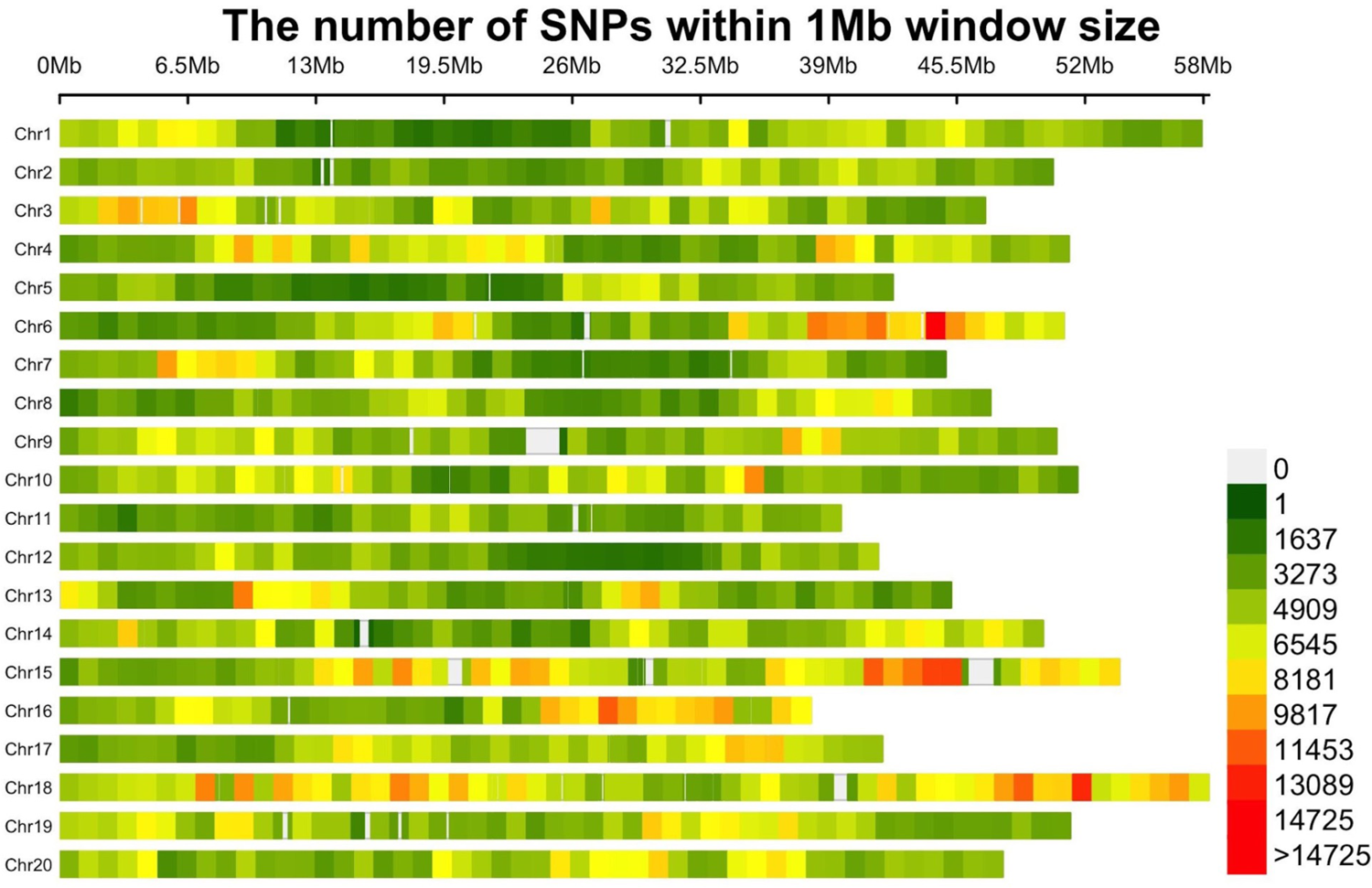
A Heatmap showcasing the distribution of Single Nucleotide Polymorphisms (SNPs) across chromosomes 1 to 20. The color gradation, from green indicating fewer SNPs to red indicating higher counts, represents the density of SNPs within 1 Mb window sizes. These SNPs are sourced from the VCF file produced through whole genome sequencing.

**Supplementary Fig. 6:**
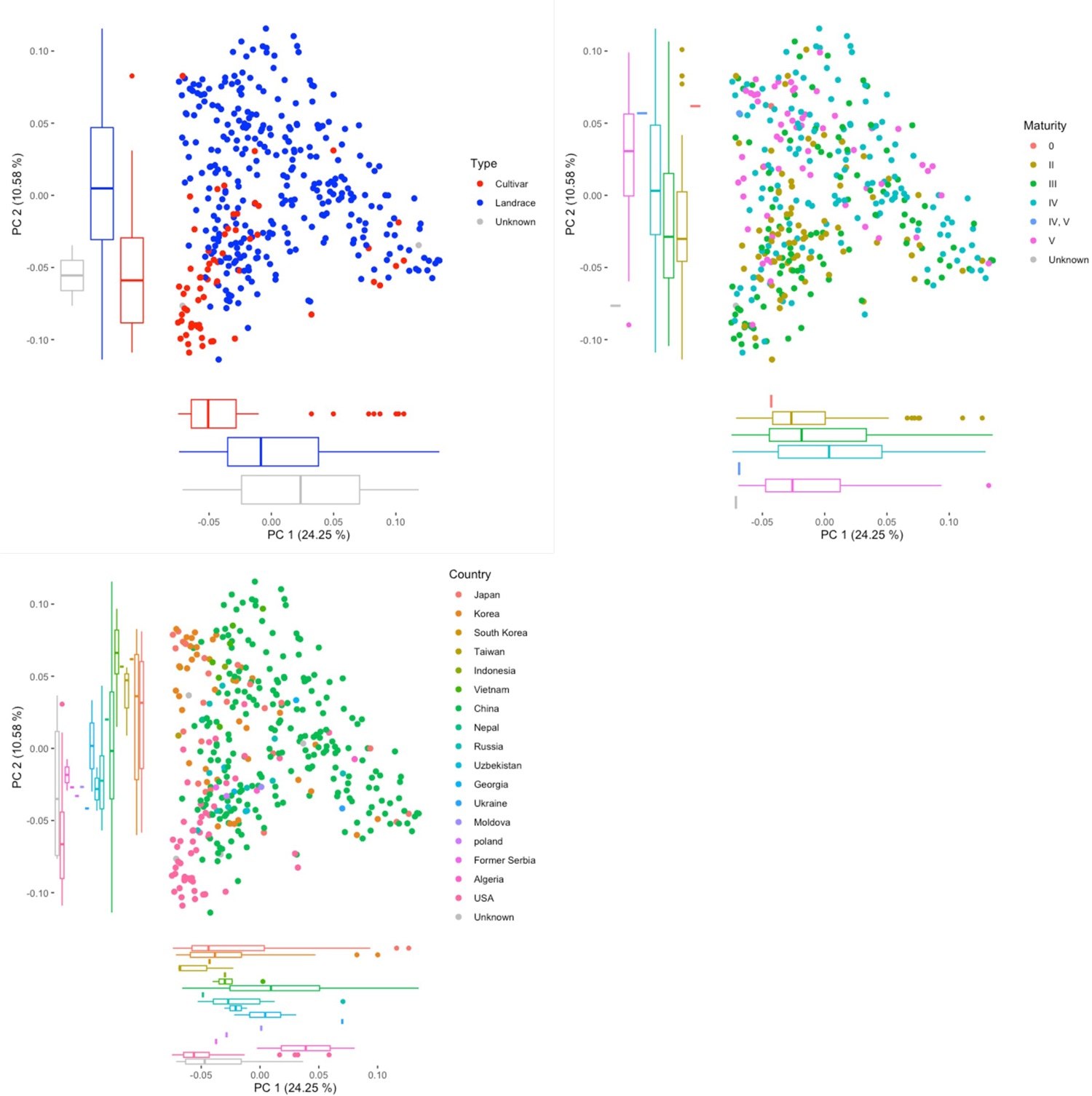
Principal Component analysis (PCA) results obtained from single nucleotide polymorphism (SNP) variations observed across 371 accessions. The box plots showcased in the figure illustrate the sample distribution, with each color denoting a distinct subgroup within the population. The population structure within this soy group exhibits divergence based on three factors: type (left), geography (right), and maturity group (bottom).

**Supplementary Fig. 7:**
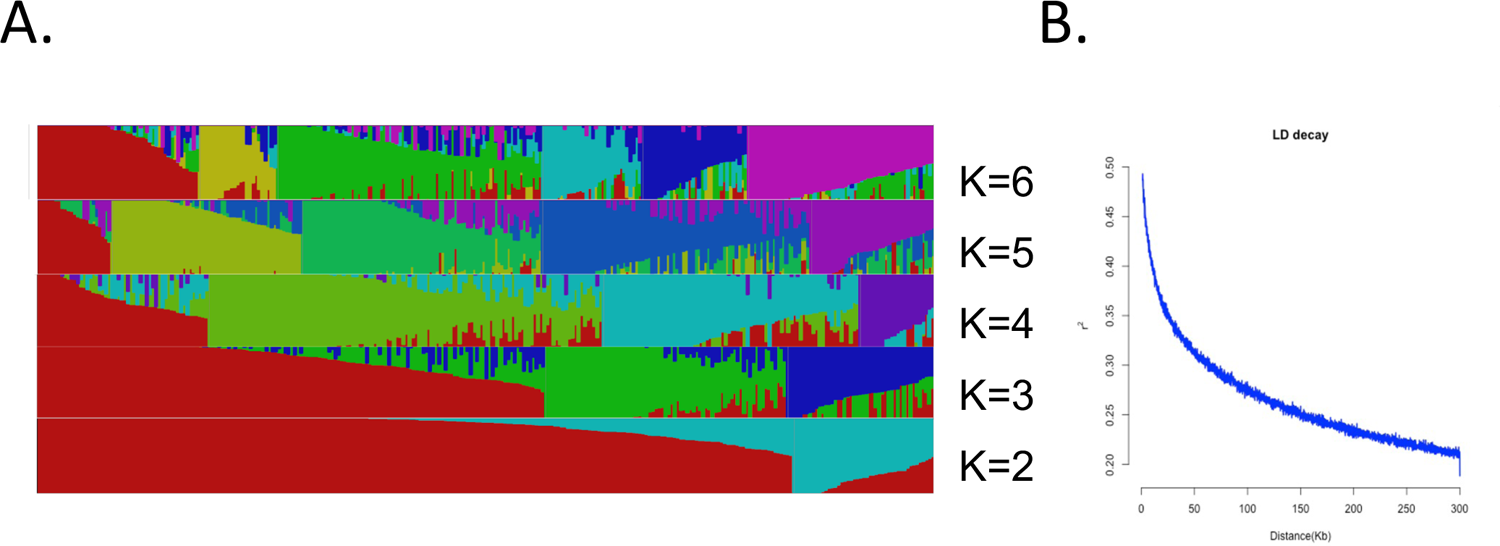
A. The population structure of soy accessions was calculated using fastSTRUCTURE, delineating distinct subpopulations based on genetic variations among the 371 soy accessions. Different population assignments are visualized by different colors. B. The mean value of linkage disequilibrium (LD) was calculated within genomic regions spanning 100 to 300 kb to elucidate patterns of LD over intermediate genomic distances. The plot provides the distribution of LD across the genome up to 300 kb.

**Supplementary Fig. 8:**
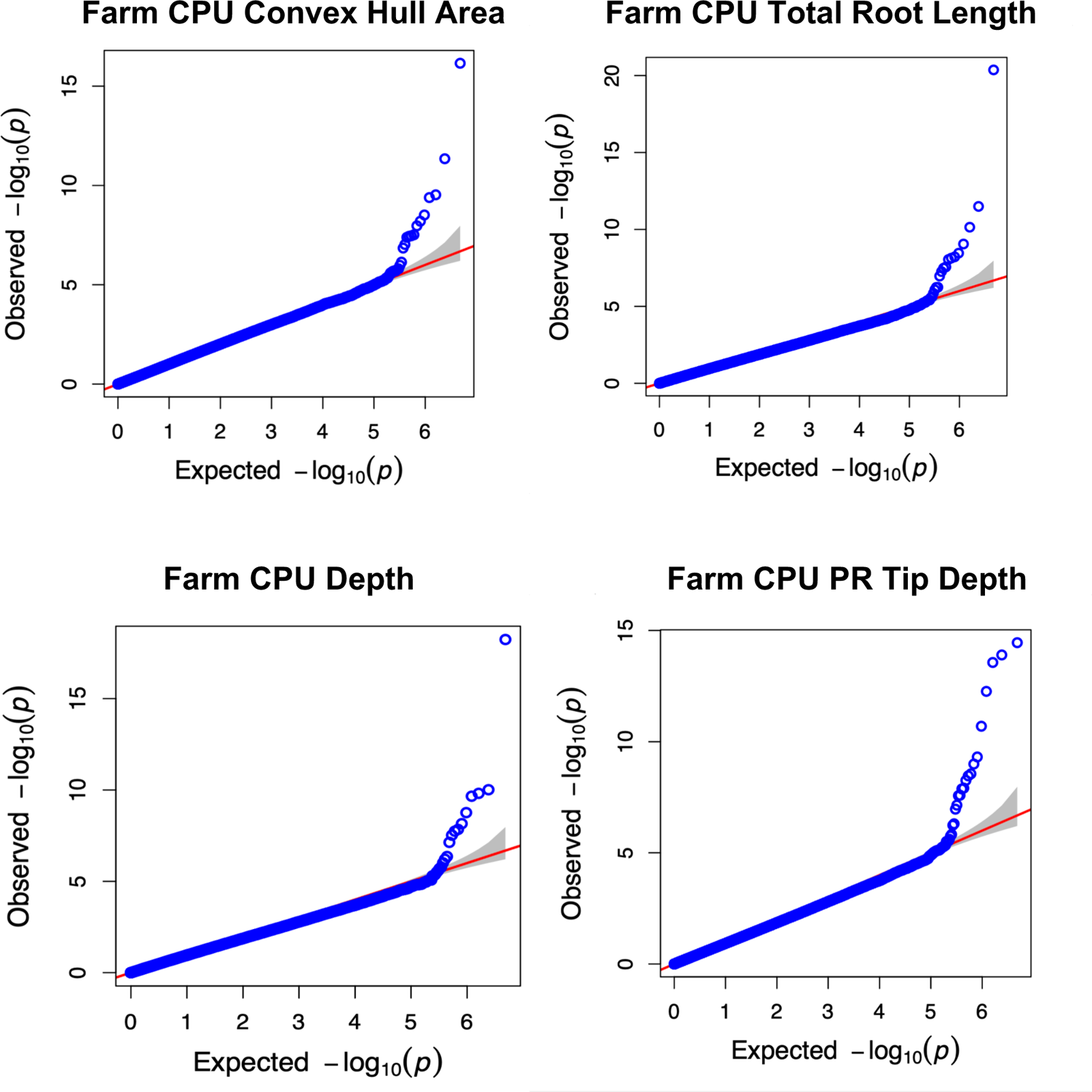
quantile-quantile (QQ) plots. The Y-axis is the observed negative base 10 logarithm of the P-values, and the X-axis is the expected observed negative base 10 logarithm of the P-values under the assumption that the P-values follow a uniform distribution. The red line and grey banding show the 95% confidence interval for the QQ-plot under the null hypothesis of no association between the SNP and the trait. Blue circles are the observed–expected P-values, which show no evidence for systematic spurious associations. Only a subset of high P-values are plotted.

**Supplementary Fig. 10:**
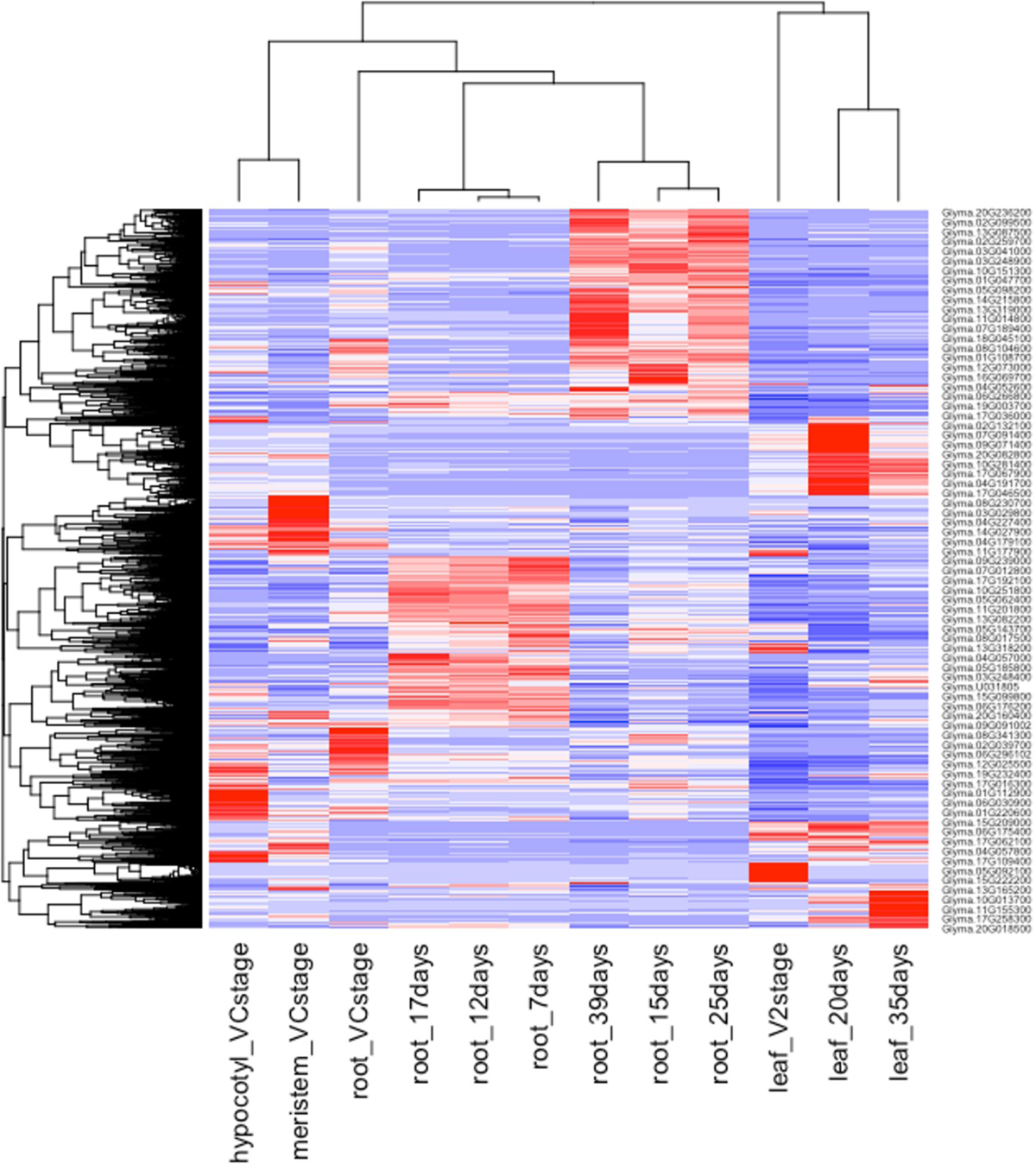
Heatmap highlights differentially expressed genes (DEGs) enriched in specific tissues and developmental stages, with red indicating high expression and blue signifying low expression.

**Supplementary Figure 11:**
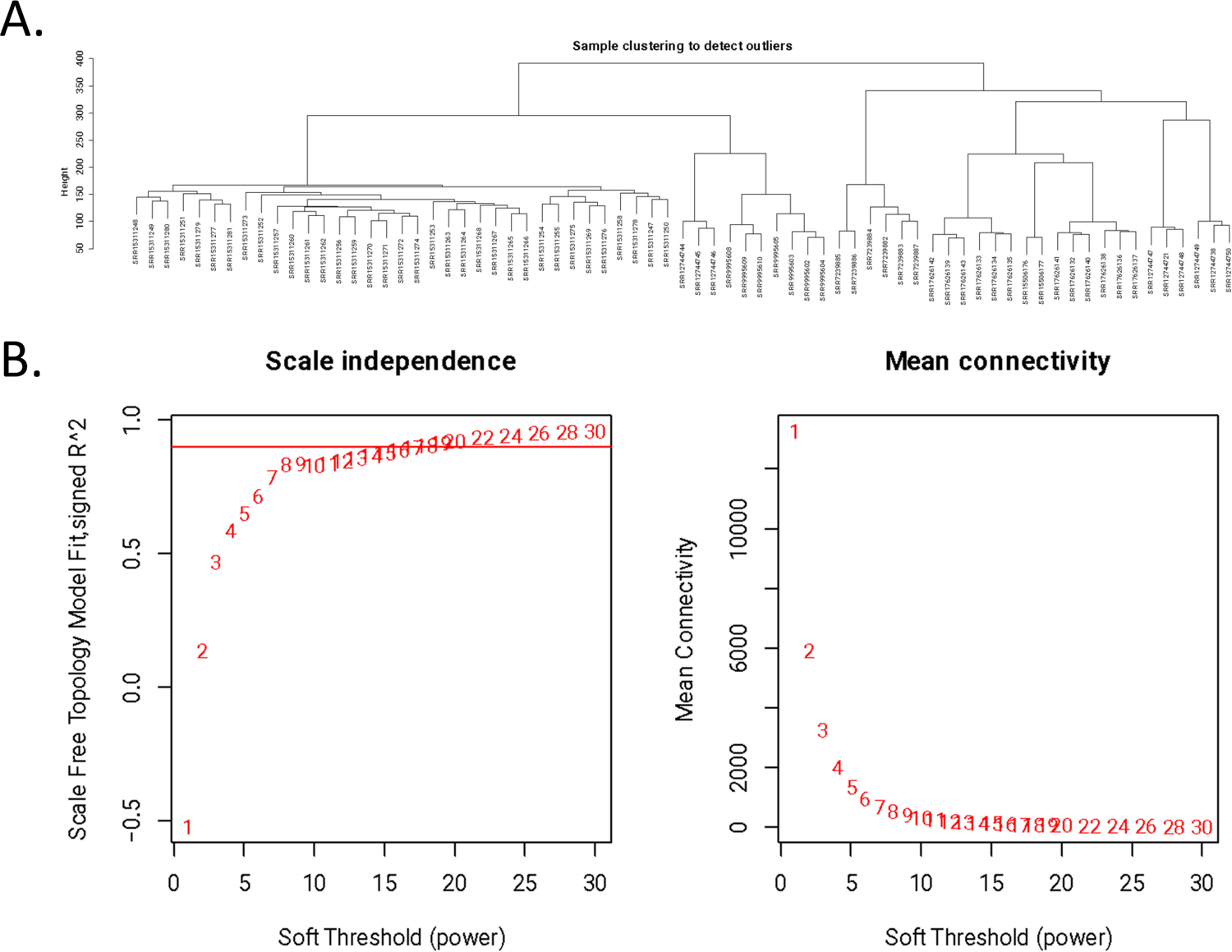
A. The clustering dendrograms of samples. B. Soft thresholding determined by the scale free network index. The left indicates the screen of optimal soft threshold. The right indicates its impact on the mean network connection level.

**Supplementary Fig. 12:**
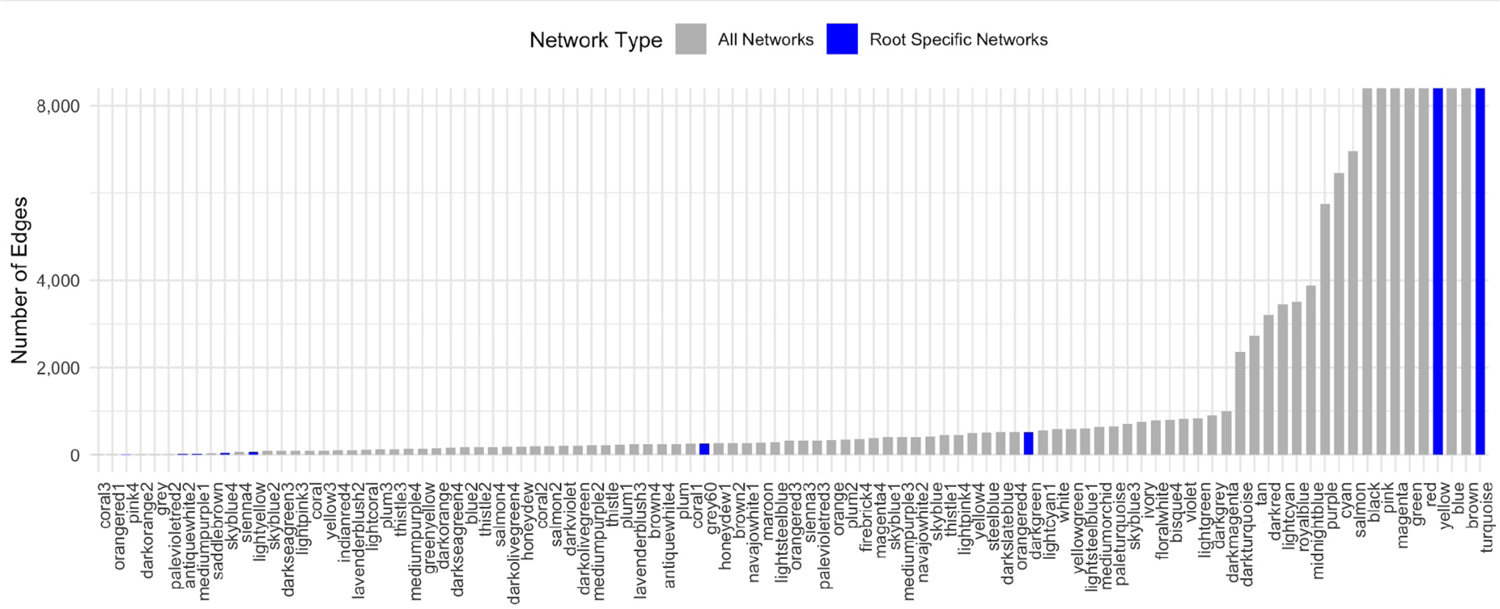
The bar chart illustrates the number of root-specific networks identified in our analysis. It highlights that the most predominant network, turquoise, had the highest count of root-specific Differentially Expressed Genes (DEGs), while several smaller modules also exhibited root-specific associations.

**Supplementary Fig. 13:**
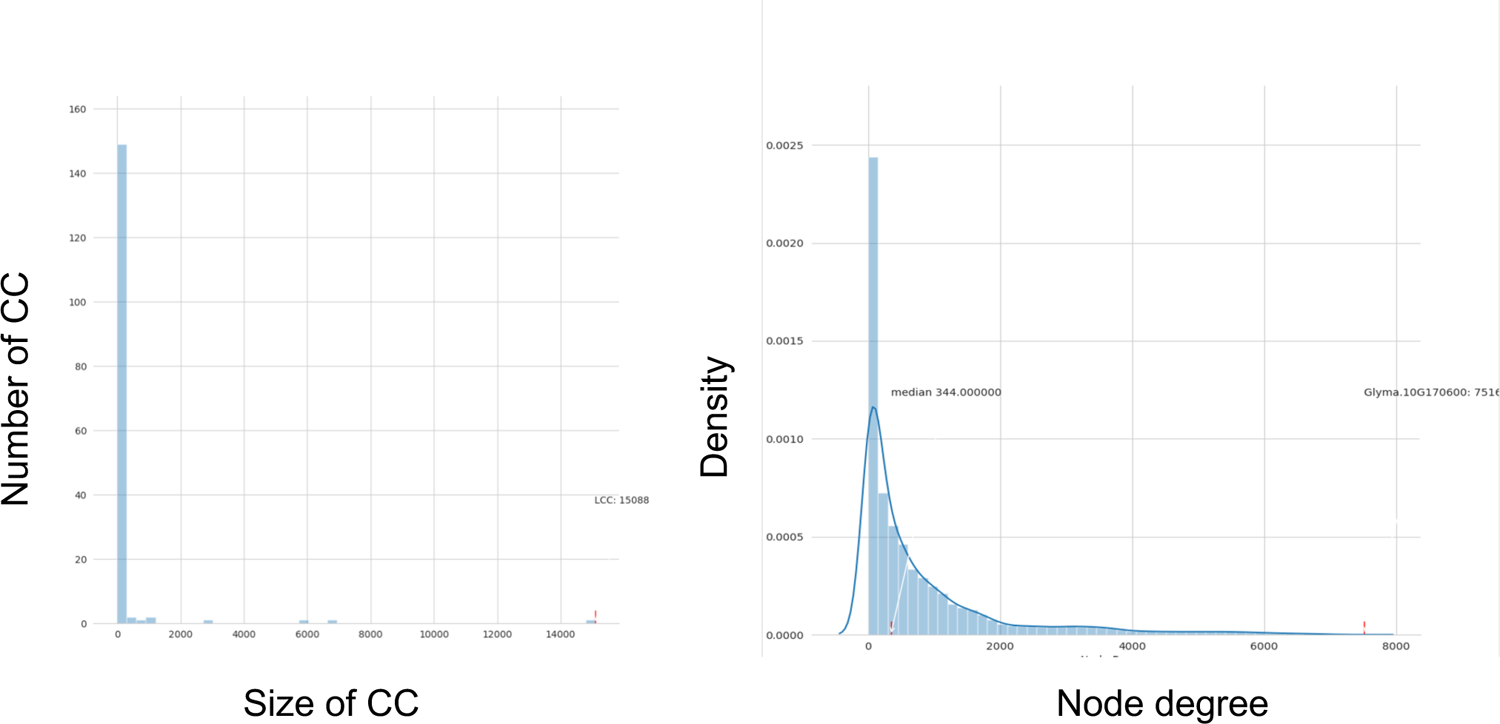
All network components where left side shows the number of networks vs the size of the networks (size of cc) and the right side shows the number of nodes with that connection vs the number of connections (node degree)

**Supplementary Fig. 14:**
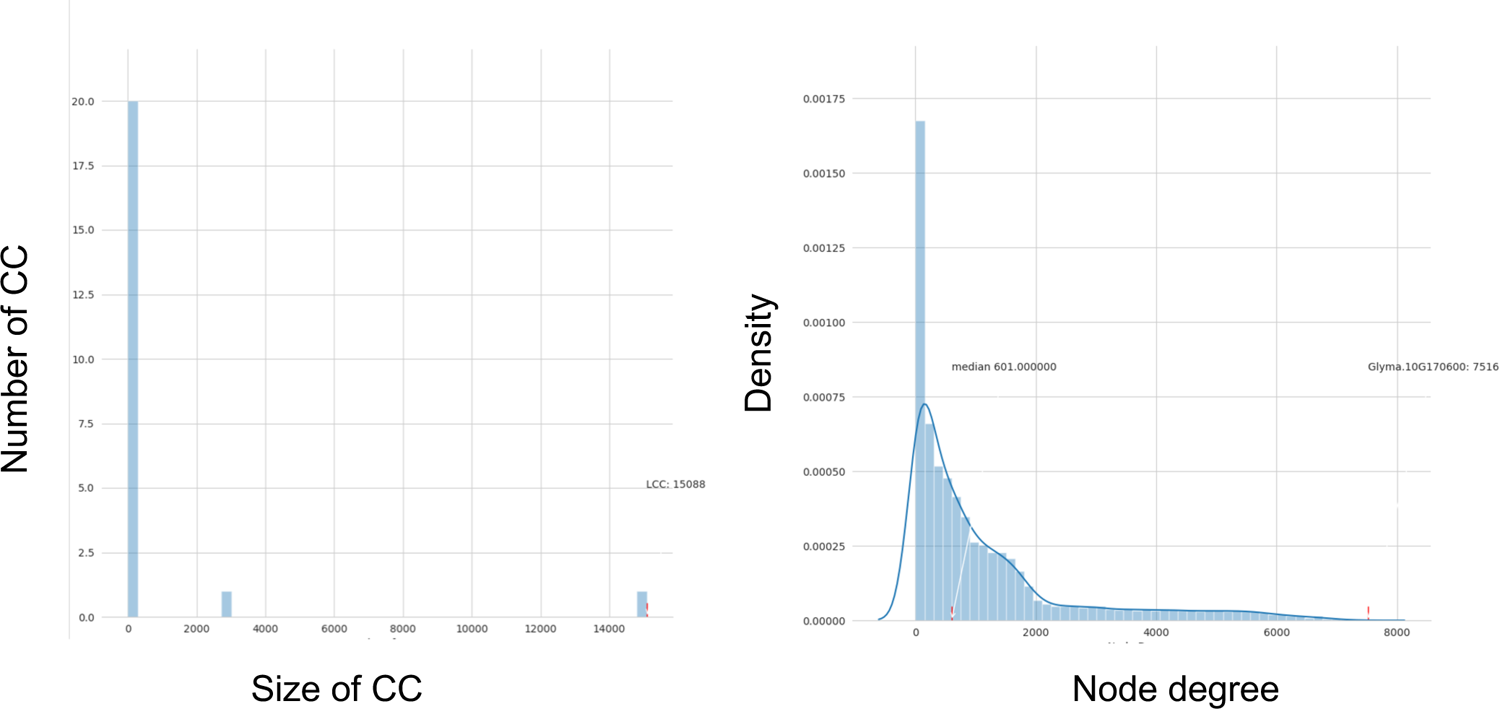
Root enriched network components where left side shows the number of networks vs the size of the networks (size of cc) and the right side shows the number of nodes with that connection vs the number of connections (node degree)

**Supplementary Fig. 15:**
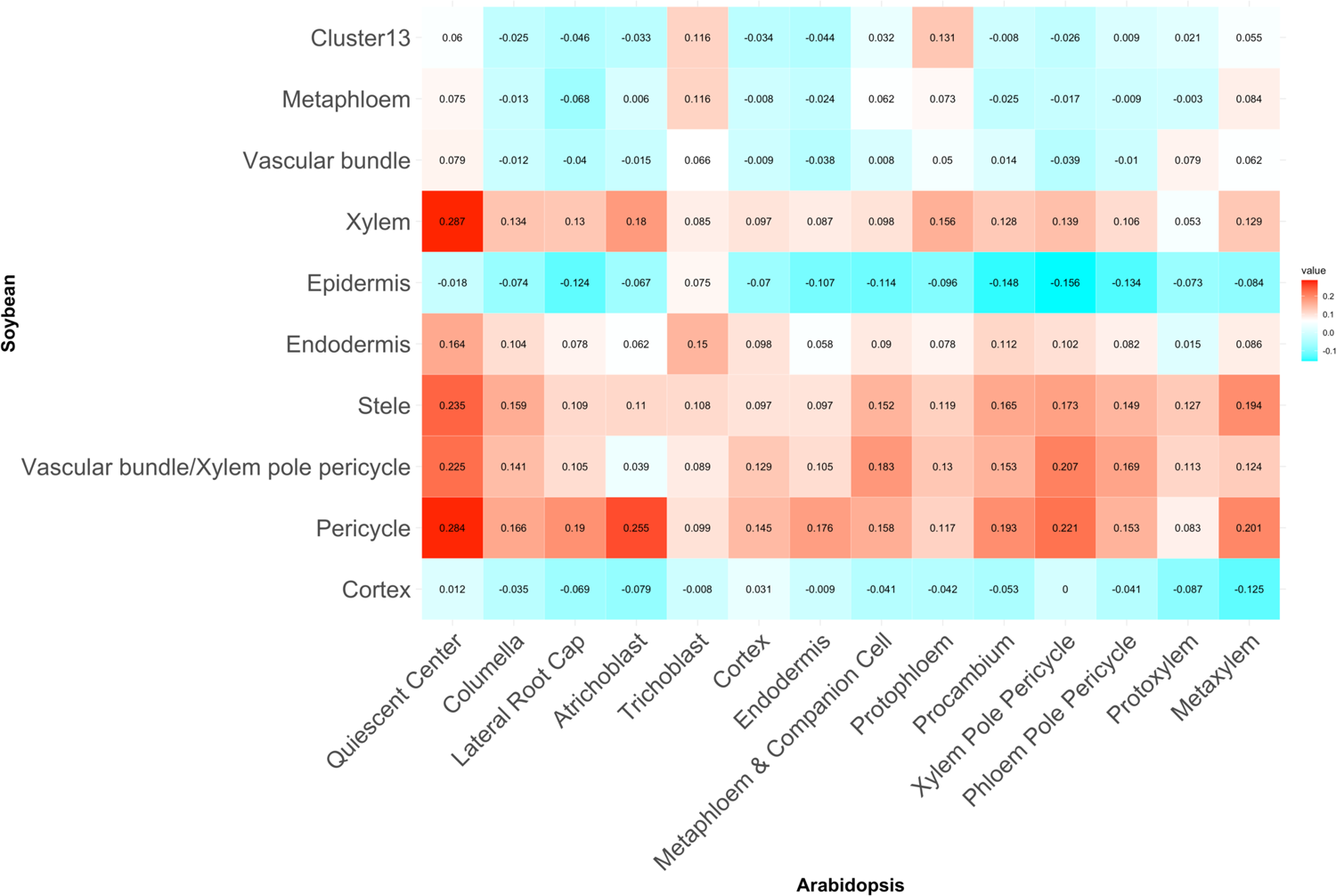
Correlation of cell types/clusters between Arabidopsis and Soybean where Spearman rank order correlation coefficient values in red indicate positive correlation and blue indicate negative correlation.

**Supplementary Fig. 16:**
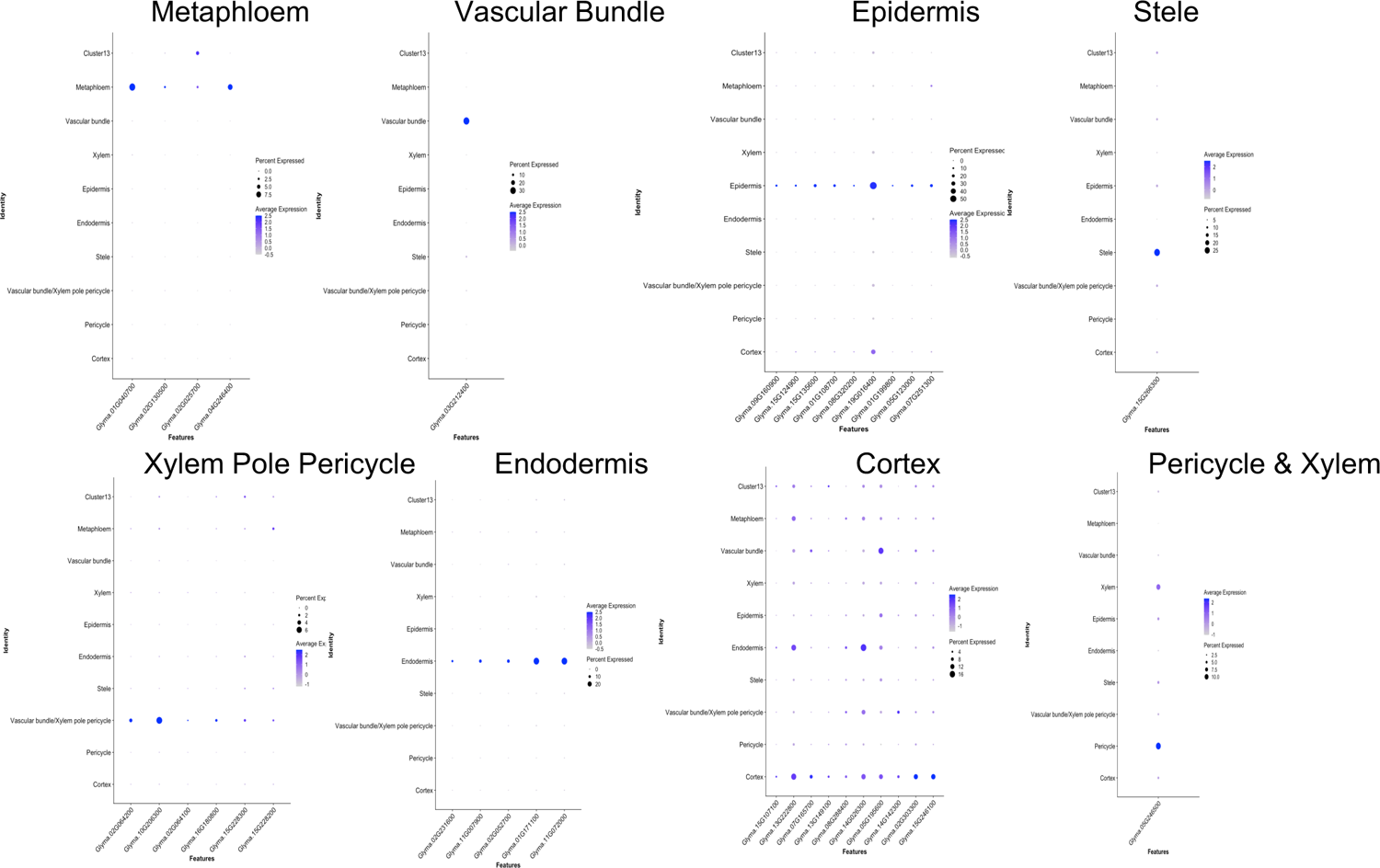
Expression pattern of cell-type specific marker genes. Dot plots indicate marker genes for metephloem, vascular bundle, epidermis, stele, xylem pole pericycle, endodermis, cortex, and pericycle & xylem.

**Supplementary Fig. 17:**
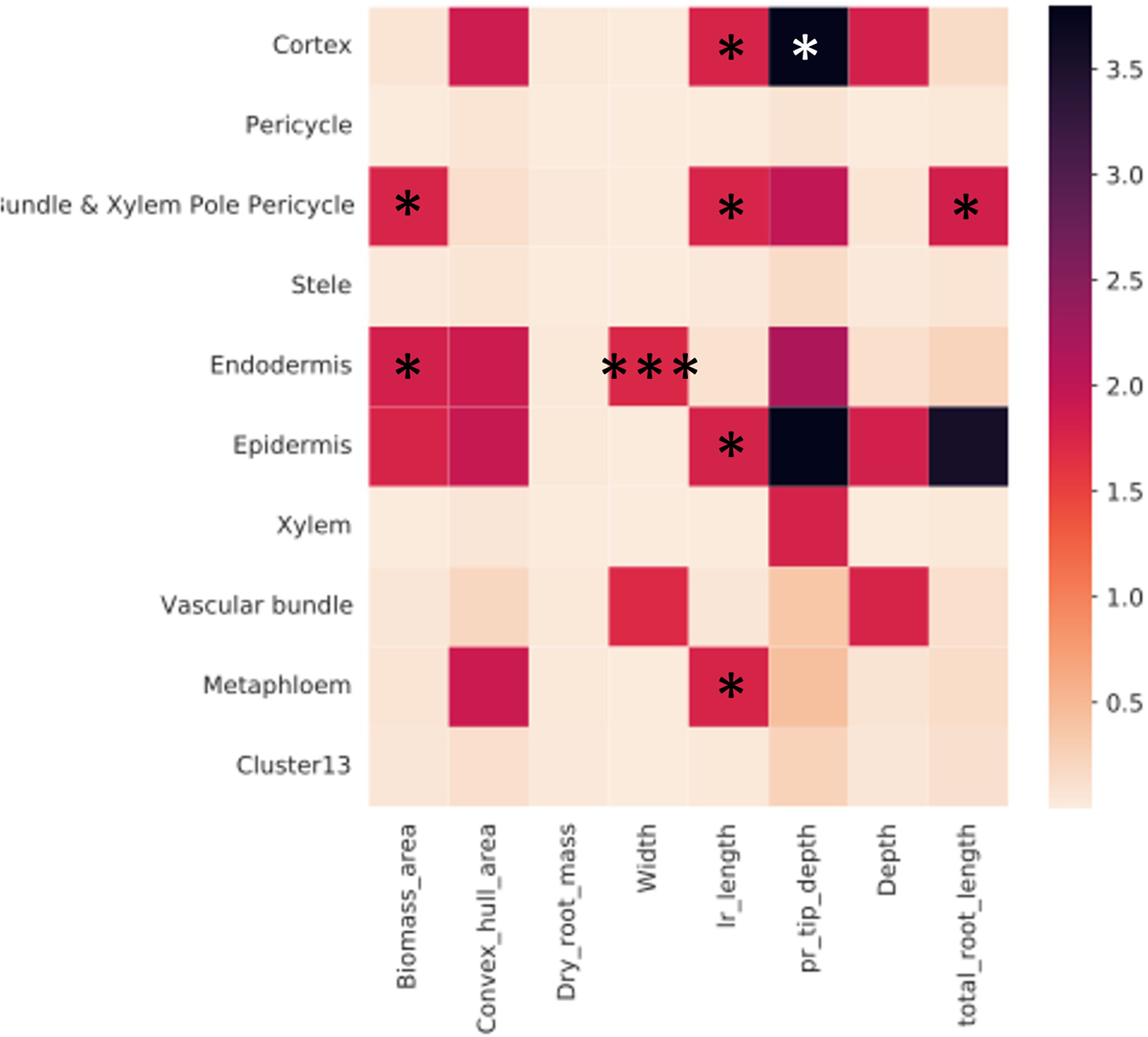
Heatmap indicating significant results from the Random Walk with Restart test from PYGNA. Read and black colors indicate that the observed is significantly far away from the null. * = p-value <0.01, ** = p-value <0.005, *** = p-value <0.001.

**Supplementary Fig. 18:**
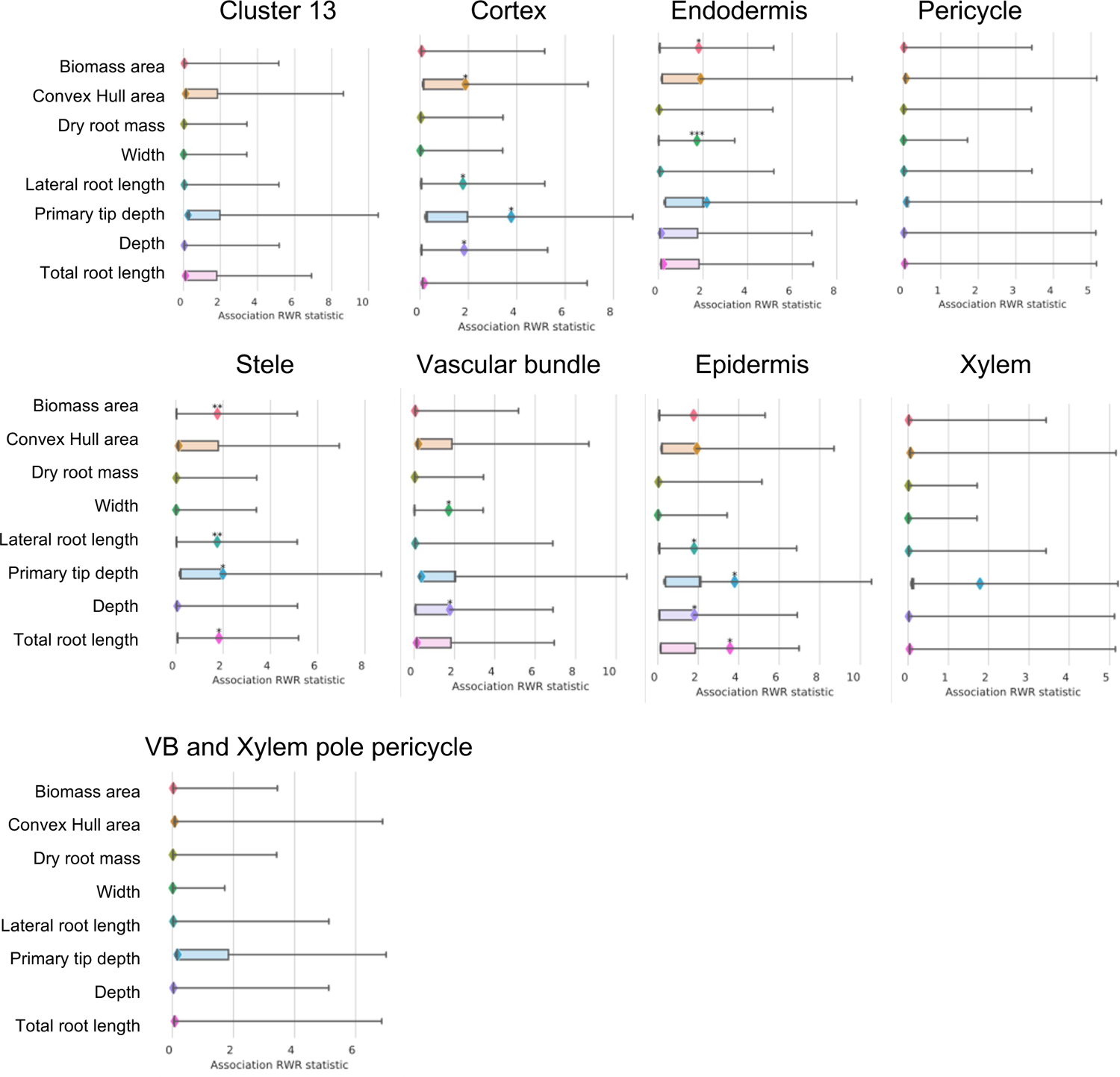
Gene set analysis of different cell types with GRIT genes separated by trait to identify subnetworks within the root GCN. The analysis was performed using the Random Walk with Restart test from PYGNA. The boxes represent null distributions, and the whiskers display the entire range. The observed values are indicated by diamonds.

**Supplementary Fig. 19:**
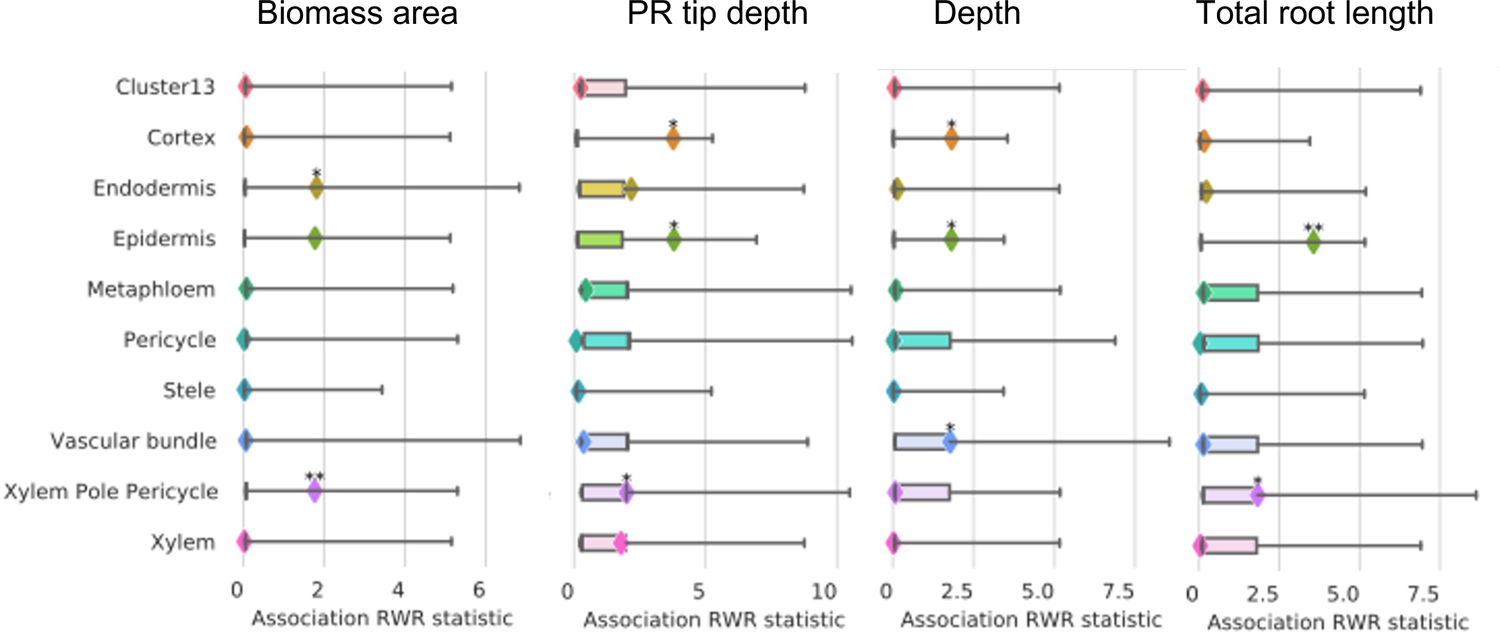
Gene set analysis of GRIT genes expressed in the single-cell data to identify subnetworks within the root GCN. The analysis was performed using the Random Walk with Restart test from PYGNA. The boxes represent null distributions, and the whiskers display the entire range. The observed values are indicated by diamonds.

**Supplementary Fig. 20:**
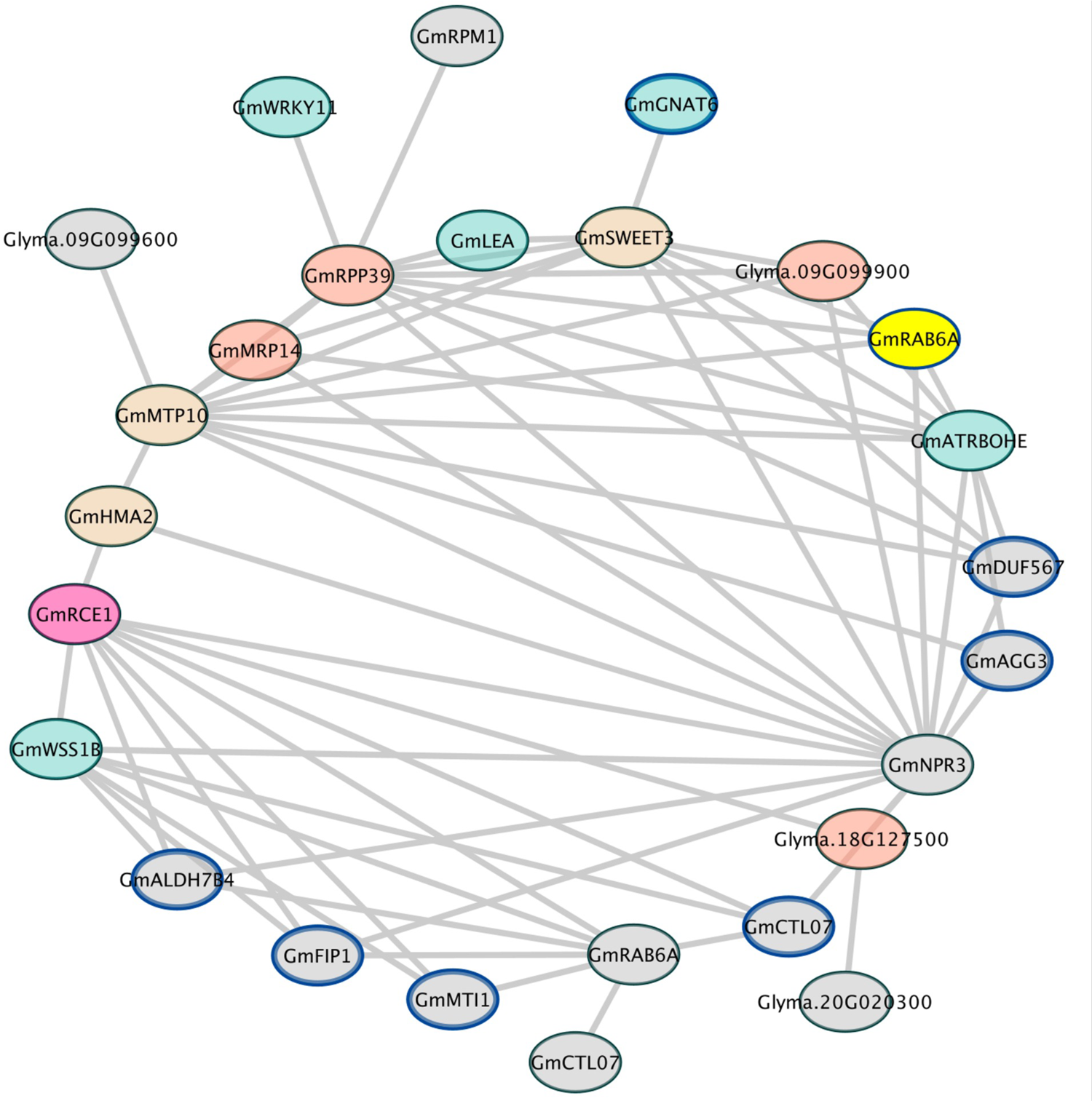
The network illustrates the connection between genes associated with lateral root length (marked with a blue border) and other genes identified through Genome-Wide Association Studies (GWAS). Different colors represent specific cell types or tissues, such as metaphloem in red, XPP in tan, epidermis in blue, and cortex in orange. Additionally, GWAS genes are denoted by the color grey.

## Notes

https://gitlab.com/salk-tm/soybean-root-gwas/

https://github.com/Salk-Harnessing-Plants-Initiative/SSRAPC-Soy-Segmentation-Root-Architecture-Phenotyping-for-Cylinder.git

https://pypi.org/project/pygna2/

https://gitlab.com/salk-tm/pygna2

